# Single-step Enzymatic Glycoengineering for the Construction of Antibody-cell Conjugates

**DOI:** 10.1101/279240

**Authors:** Jie Li, Mingkuan Chen, Zilei Liu, Linda Zhang, Brunie H. Felding, Gregoire Lauvau, Michael Abadier, Klaus Ley, Peng Wu

## Abstract

Employing live cells as therapeutics is a direction of future drug discovery. An easy and robust method to modify the surfaces of cells directly to incorporate novel functionalities is highly desirable. However, many current methods for cell-surface engineering interfere with cells’ endogenous properties. Here we report an enzymatic approach that enables the transfer of biomacromolecules, such as a full length IgG antibody, to the glycocalyx on the surfaces of live cells when the antibody is conjugated to the enzyme’s natural donor substrate GDP-fucose. This method is fast and biocompatible with little interference to cells’ endogenous functions. We applied this method to construct two antibody-cell conjugates (ACCs) using different immune cells, and the modified cells exhibited specific tumor targeting and resistance to inhibitory signals produced by tumor cells, respectively. Remarkably, Herceptin-NK-92MI conjugates exhibits enhanced activities to induce the lysis of HER2+ cancer cells both ex vivo and in a murine tumor model, indicating its potential for further development as a clinical candidate.

---

With cell engineers’ tireless efforts of converting cells into living therapeutics over the past decade, spectacular results have been observed in patients treated with adoptive cell transfer (ACT)^1^. The most remarkable example is *Kymriah*, a chimeric antigen receptor T-cell (CAR-T) therapy that was approved recently as the first cell-based gene therapy in the United States^2^.

The major technical challenge in cell engineering is to confer new properties to the manipulated cells with little interference with the cells’ endogenous functions. As the most common and robust cell-engineering approach, genetic engineering is limited by technical complications and safety concerns (**Fig. 1A**), such as the viral transduction resistance of primary cells, heterogeneous expression levels, and the potential for endogenous gene disruption^3-5^. Therefore, engineering cell surfaces from “outside” has emerged as a complementary and generally-applicable approach^6,7^. Preeminent examples include metabolic oligosaccharide engineering (MOE) developed by Bertozzi, *et al.* (**Fig. 1A**) and sortagging, the transpeptidation reaction catalyzed by bacterial sortases, among others^6,8-10^. MOE requires a two-step procedure, combining metabolic labeling with bioorthogonal chemistry to endow cell-surface glycans with new functions. Sortagging involves only a single-step treatment. However, cells without genetic modification only have a few thousand naturally exposed glycine residues that can be functionalized by this approach^11^. Therefore, direct functionalization of the cell surface in a non-invasive and highly efficient way is still difficult to achieve^6,10^. Cell engineering would benefit from a single-step method that efficiently modifies native substrates on the surfaces of cells to incorporate novel functionalities.

**Figure 1.**
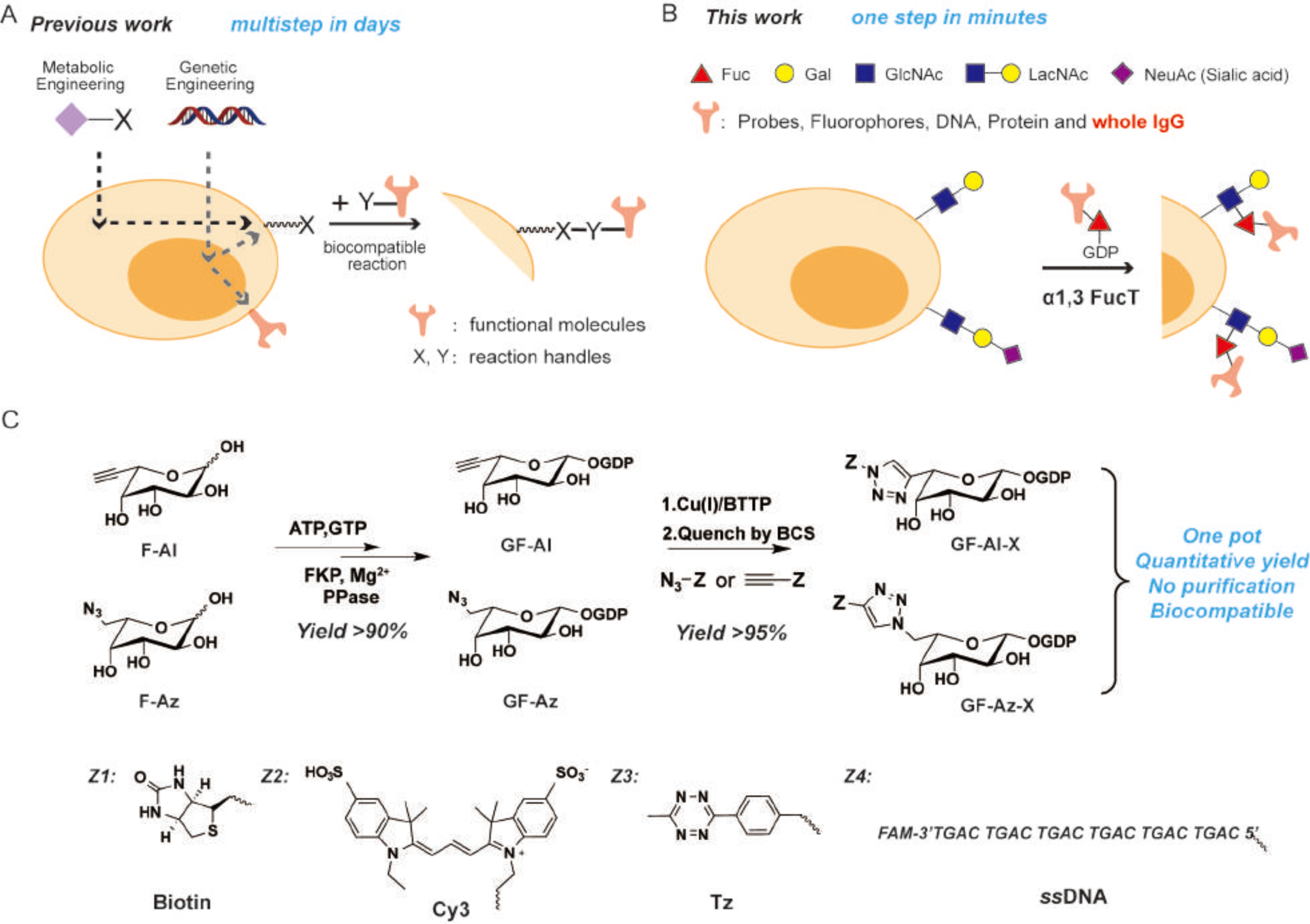
The One-step fucosylation-based strategy for cell-surface engineering: (**A**) Two representative cell-surface engineering approaches. Metabolic engineering is used to install a reaction handle (X) onto the surface of the cell, which can react with a complementary handle (Y) on a molecule of interest. Genetic engineering allows the direct expression of functional molecules or the installation of reaction handles (X) on the surfaces of cells. (**B**) This work reports on an enzymatic glycoengineering approach capable of transferring a variety of functional molecules to the surfaces of cells in one step. The reaction between LacNAc/sialylLacNAc and GDP-Fuc derivatives on the surfaces of cells is enabled by *H. pylori* α1,3FucT that tolerates modifications as large as a whole IgG conjugated at the C6 position of fucose; (**C**) the one-pot protocol for the synthesis of GF-Al and GF-Az derivatives. The new functional group (Z) conjugated to fucose includes bioorthogonal handles (tetrazine, Tz), biophysical probes (biotin, Cy3), and biomaterials (*ss*DNA).

*In situ* glycan editing via glycosylation enzymes is a single-step approach to modify glycocalyx. The most notable example of its application is *ex vivo* fucosylation of mesenchymal stem cells and regulatory T cells using GDP-fucose (GDP-Fuc or GF) and recombinant human α(1,3)-fucosyltransferase (FucT or FT) VI ^12,13^. This procedure, currently undergoing several clinical trials, improves adhesion, homing, and engraftment of adoptively-transferred cells. However, enzymatic glycoengineering has not been widely used in therapeutic interventions^6^. A major limitation is that current enzymatic transferrable substrates are confined to small, synthetic molecules (MW <2000)^14-16^, while biomacromolecules (*e.g.* monoclonal antibodies, mAbs) that have high therapeutic value are not accessible.

Here, we report the discovery of the remarkable substrate tolerance of *H. pylori* α1,3FucT. This enzyme enables quantitative transfer of a full length IgG antibody conjugated to the GDP-Fuc donor to LacNAc (Galβ1,4GlcNAc), a common building block of glycocalyx, on the cell surface of live cells within a few minutes (**Fig. 1B**). A one-pot protocol that couples the synthesis of an unnatural GDP-Fuc derivative to the subsequent derivative transfer was developed and made this engineering approach practical and cost effective. Using this technique, we constructed two types of antibody-cell conjugates (ACCs) using a natural killer cell line (NK-92MI) and primary CD8+ OT-1 T cells. We demonstrated, for the first time, the application of this technique to boost the activities of modified immune cells, including specific tumor targeting and resistance to inhibitory signals produced by tumor cells.

## Results and discussion

### A one-pot protocol for preparing and transferring GDP-Fuc derivatives

To develop the enzyme-based glycan modification as a general method for cell-surface engineering, a practical and scalable approach for the preparation and transfer of nucleotide sugar donors equipped with new functional groups is required^17^. We discovered that GDP-L-6-ethynylfucose (GF-Al) or GDP-L-6-azidofucose (GF-Az) produced *in situ* can be coupled directly with a wide variety of probes using the ligand accelerated copper(I)-catalyzed alkyne-azide cycloaddition (CuAAC)^18-20^ (**Fig. 1C**). These probes include biotin, a fluorescent probe Cy3, a bioorthogonal reaction handle tetrazine (Tz) and a dye (fluorescein amidite, FAM) labeled, single-strand DNA (*ss*DNA) (**Supplementary Fig. S1**). All reactions attained near quantitative yields (> 90%), and the crude products were rendered biocompatible for direct transfer by α1,3 FucT onto the cell surface after quenching the reaction with the FDA-approved copper chelator bathocuproine sulphonate (BCS) (**Supplementary Fig. S2**). Compared to the conventional two-step labeling protocol^15^, i.e. enzymatic transfer followed by cell surface click chemistry, the one-step enzymatic labeling using the substrate of one-pot product was significantly more efficient and biocompatible (**Supplementary Fig. S3**). In addition, we found that the enzymatic transfer of the one-pot Tz derivative made from GF-Az was more efficient than that made from GF-Al (**Supplementary Fig. S2D**).

Sortagging is probably the best known enzymatic covalent ligation reaction without the need for genetic manipulation of the target cell population. We directly compared the efficiency of the FucT-mediated cell-surface modification with that catalyzed by sortase (**SrtA** 5M) using biotin-conjugated substrates. At the optimal substrate concentration (500 μM of biotin-LPETG), negligible transpeptidation reaction took place within 2 hr in the presence of 1 μM of sortase (**Supplementary Fig. S4A**). By contrast, 0.6 μM of FucT afforded robust cell-surface labeling within 2 min in the presence of 50 μM of GDP-Fuc-biotin (**Supplementary Fig. S4B**). Even at 20 μM enzyme concentration, it took 120 min for sortagging to reach signal saturation^21^. Moreover, when 0.6 μM of FucT and 20 μM sortase were used for cell-surface modification, respectively, the FucT-catalyzed process was found to be at least 80 times more efficient than the sortase-catalyzed process (**Supplementary Fig. S4A**).

### Enzymatic transfer of full-length IgG molecules to the cell surface using FucT

The remarkable efficiency of fucosylation to transfer small-molecule probes with diverse structures to the cell surface combined with the previous known plasticity of mammalian sialyltransferases for protein PEGylation^22^ suggests the possibility that FucT may tolerate even larger molecules conjugated to the fucose C6 position. To assess this possibility, we conjugated GDP-fucose with monoclonal antibodies (mAbs, full length IgG), the fastest growing class of protein drugs. The bioorthogonal handle *trans*-cyclooctene (TCO) with a PEG linker was installed onto mAbs or their isotype controls via standard amine-coupling procedures^23^. Subsequently, mAbs bearing TCO moieties were reacted with GF-Az-Tz via the inverse electron-demand Diels–Alder reaction (IEDDA)^24^ to generate GDP-Fuc-conjugated IgG molecules (GF-IgG) (**Fig. 2A**). GDP-Fuc modified antibodies were characterized by MALDI-TOF MS and were found to exhibit similar antigen binding capacities compared to their parent antibodies (**Supplementary Fig. S5**).

**Figure 2.**
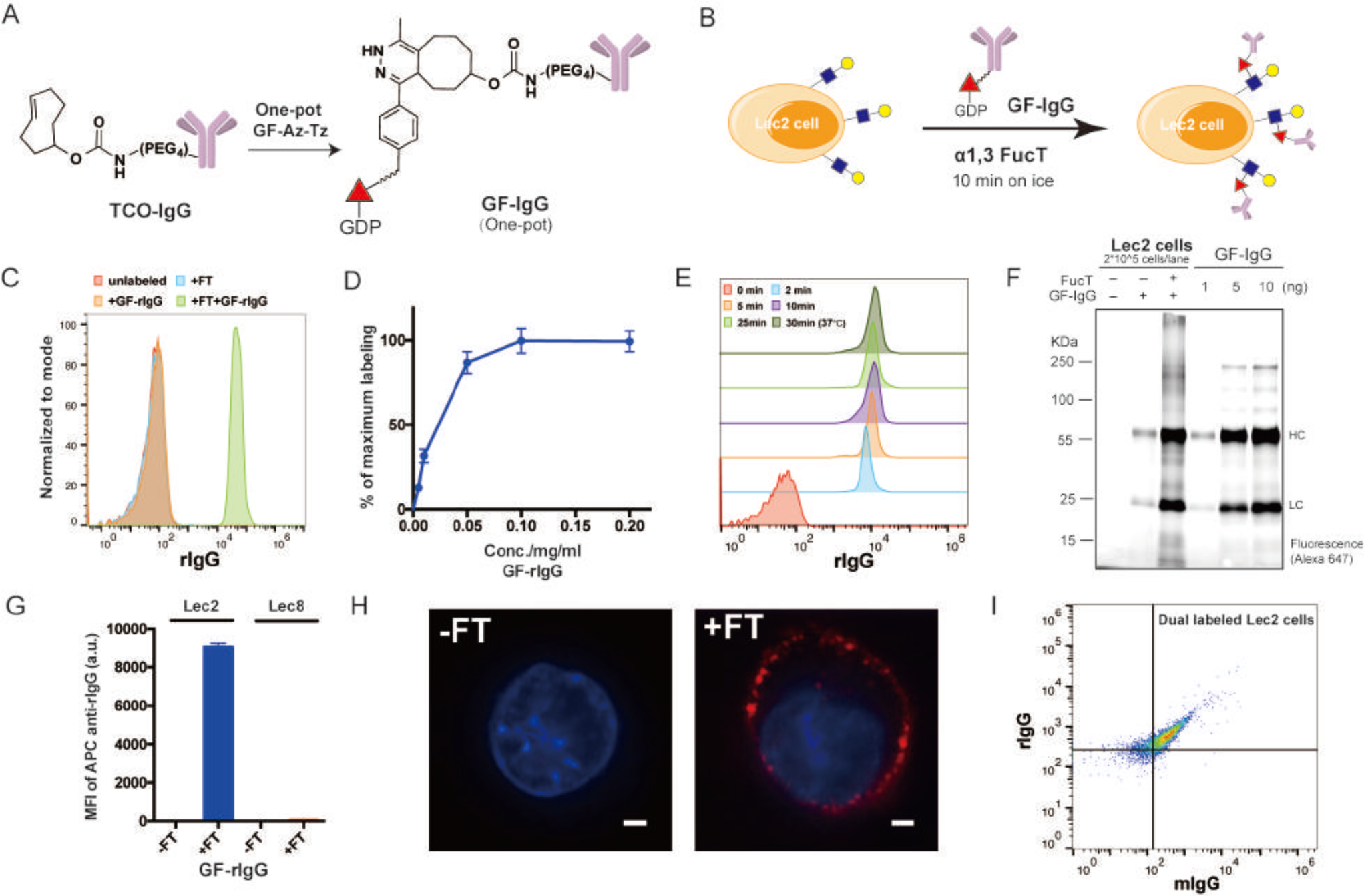
Enzymatic transfer of IgG to the surfaces of Lec2 CHO cells: (**A**) Schematic representation of the synthesis of a GDP-Fuc conjugated IgG (GF-IgG). (**B**) Workflow of the FucT-catalyzed transfer of GF-IgG to the surface of Lec2 CHO cells. (**C**) Flow cytometry analysis of Lec2 cells treated with enzyme FucT, substrates GF-rIgG, or both. (**D**) Titration of GF-rIgG, concentrations ranging from 0.005 mg/ml to 0.2 mg/ml in the reaction buffer; each reaction used the same amount of FT and proceeded at room temperature for 30 min; Mean ± SD (error bars), representative graph from three independent experiments; (**E**) Time course of enzymatic transfer of GF-rIgG to Lec2 cells on ice; reaction at 37 °C was used as the maximum labeling control; (**F**) Fluorescent gel imaging of detecting and quantifying rIgG (Alexa Fluor 647 labeled) molecules conjugated on Lec2 cell surface; (**G**) Lec8 CHO cells without LacNAc expression were compared with Lec2 cells in the enzymatic IgG transfer as a negative control; Mean ± SD (error bars), representative graph from three independent experiments; (**H**) Confocal microscopy images of Lec2 cells treated with or without FT when incubated with Alexa Fluor 647 labeled GF-rIgG; nuclei were stained with Hoechst 33342. Scale bar: 2 μm; (**I**) Flow cytometry analysis of Lec2 cells simultaneously labeled with rIgG and mIgG.

The one-pot product of GDP-Fuc conjugated rat IgG (GF-rIgG) was then incubated with Lec2 CHO cells that express abundant terminal LacNAc units in the presence of FucT (**Fig. 2B**). Remarkably, the signal of rIgG conjugated onto the cell surface was detectable after a 2-min incubation with FucT (60 mU) and GF-rIgG (0.1 mg/ml) (**Fig. 2C**). The labeling efficiency was concentration-dependent (GF-rIgG), which reached saturation at 0.1 mg/ml (**Fig. 2D**). Notably, the conjugation reaction was completed in 10 minutes even on ice (**Fig. 2E**). At the saturated condition, approximately 2.5×10^5^ rIgG molecules were introduced to the cell surface (**Fig. 2F and Supplementary Fig. S6A**). The viability of the rIgG-labeled cells was similar to that of unlabeled cells, which further confirmed the biocompatibility of this one-pot procedure (**Supplementary Fig. S7**). Lec8 CHO cells, which do not express LacNAc, were used as a negative control to confirm that the transfer was dependent on the reaction between the GDP-Fuc derivative and cell-surface LacNAc. As expected, only background fluorescence was displayed by the Lec8 cells after the enzymatic reaction (**Fig. 2G**). A competition experiment using cells blocked by natural substrates of FucT, i.e., GDP-Fuc, also was conducted, and the subsequent GF-rIgG labeling almost was abolished (**Supplementary Fig. S8**). These results confirmed that the reaction sites of enzymatic transfer of GF-rIgG were the same as those for GDP-Fuc on the surface of the cell. Furthermore, confocal microscopy analysis verified that most of the labeled rIgGs were located on the cell membrane (**Fig. 2H**). Importantly, multiple functionalities can be introduced to the surface of the cell simultaneously, e.g., two antibodies (GF-rIgG and GF-mIgG—GDP-Fuc modified mouse IgG) (**Fig. 2I and Supplementary Fig. S9**). Such results are difficult to achieve via genetic approaches. Worthy of note, ACCs can also be built using recombinant sialyltransferases and CMP-Sialic acid-antibody conjugates though with much less efficiency (**Supplementary Fig. S10**).

This approach was equally efficient to modify primary human cells, e.g. T cells; robust labeling with IgGs was achieved within 15 min (**Supplementary Fig. S11 and S6B**). We confirmed that the bioconjugation of IgG molecules to the cell surface had no short-term interference with the expression of cell surface markers (**Supplementary Fig. S12**). The half-life of IgG molecules conjugated to the cell surface is approximately 24 hours, and the conjugation had no effect on the proliferation of the modified cells (**Supplementary Figs. S11C and S11D**).

Taken together, we confirmed that the transfer of GF-IgG to LacNAc on the cell surface via FucT is a highly efficient one-step approach to construct ACC. With this powerful method in hand, we explored its application to construct ACCs using various immune cells for boosting the efficacy of cell-based therapies.

### Herceptin-NK-92MI conjugates enables specific killing of Her2+ tumor cells in a murine model

Specific targeting is key for the success of cell-based cancer immunotherapy. In innate immunity human natural killer (NK) cells play crucial roles in the host-rejection of tumors and virally infected cells^25^. NK-92, a constantly active and non-immunogenic natural killer (NK) cell line, is being developed in bulk quantities under GMP conditions as an “off-the-shelf therapeutic” for adoptive NK-based cancer immunotherapy^25,26^. However, NK-92 cells do not express Fc receptors for antibody-dependent cell-mediated cytotoxicity (ADCC), a mechanism for specific cell lysis, which significantly limits their clinical applications^25^. We speculate that modifying NK-92 cells with antibodies against tumor specific antigens via the chemoenzymatic conjugation may confer NK-92 cells with specific targeting capability (**Fig. 3A**). We chose, NK-92MI cells, an IL-2 independent variant of the NK-92 cell line, as the candidate for bioconjugation with Herceptin because it constantly expresses a high level of LacNAc to be modified by GDP-Fuc conjugated human IgG (GF-hIgG) (**Supplementary Fig. 13A**). Herceptin, also known as Trastuzumab, is used extensively to treat human epidermal growth factor receptor 2-positive (HER2+) breast cancer. We calculated that approximately 3×10^5^ Herceptin molecules were conjugated to the surface of a NK-92MI cell, which equaled to ∼7.5 ng Herceptin/0.1 million cell (**Supplementary Fig. S6C**). Herceptin conjugated to the surface of NK-92MI cells maintains exclusive binding to the HER2 antigen (**Fig. 3B and Supplementary Figs. 13B and 13C**), and its cell-surface half-life is approximately 20 h (**Supplementary Fig. 13D**). As revealed by flow cytometry analysis and confocal imaging NK-92MI cells conjugated with Herceptin had strong interactions with BT474—a HER2+ breast cancer cell— in a co-culture assay, whereas unmodified NK-92MI cells were only weakly bound (**Figs. 3C and 3D**). Moreover, NK-92MI cells modified with Herceptin induced the lysis of BT474 cells more effectively than unmodified NK-92MI cells (**Fig. 3E**). Neither isotype control hIgG labeling nor co-treatment with excess free Herceptin (1000 ng/0.1 million NK cells) could enhance the killing activity of NK-92MI on BT474, indicating that covalent conjugation of Herceptin to the surface of NK-92MI cells is required (**Fig. 4E**). Importantly, the cell-lysis efficiency of the modified NK-92MI cells also was dependent on the loading of GF-Herceptin on the cell surface, which reached the plateau when 0.1mg/mL of GF-Herceptin was used for bioconjugation (**Fig. 3F, Supplementary Fig. 14**). The enhanced killing effect of Herceptin-NK92-MI conjugates later was confirmed on other HER2+ cancer cells, including SKBR3 and MDA-MB-435/HER2+, but not on HER2 negative (HER2-) cancer cells, such as MDA-MB-435 and MDA-MB-468 (**Fig. 3G**). In addition, the total secretion of granzyme B increased only when Herceptin-NK-92MI conjugates was mixed with BT474 (**Supplementary Fig. 13E**), indicating that the Herceptin-HER2 binding induced proximity between the modified NK-92MI cells and target cells promoted the cancer-cell triggered NK-92MI cell activation. Similar to other cell-mediated cytotoxicity, higher effector-to-target cell ratios also have better killing results, but only in the Herceptin-labeled NK-92MI group, which reaches saturation at 5:1 (**Fig. 3H**). Another distinct advantage of enzymatic cell engineering is that several antibodies can be conjugated onto the surface of a cell at the same time. As proof-of-concept, we conjugated NK-92MI cells with both of Herceptin and an anti-EGFR antibody (**Supplementary Fig. 15**), and the duel modified cells exhibited better killing efficiency on SKOV3 cells (HER2+EGFR+) than that induced by the single-mAb modified counterparts (**Fig. 3I**).

**Figure 3.**
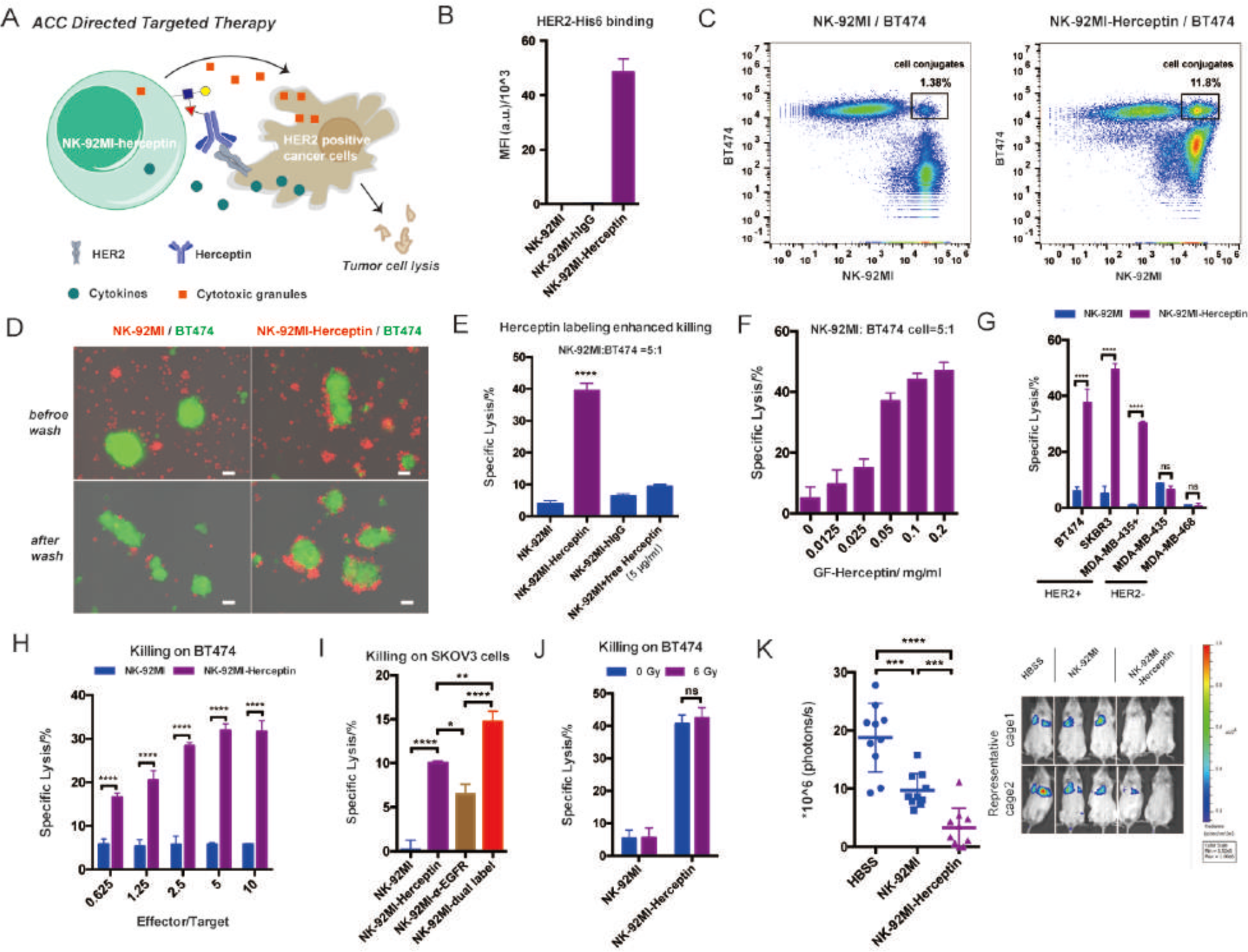
Construction of Herceptin-NK-92MI conjugates for targeting HER2+ cancer cells: (**A**) Hercetin-NK-92MI conjugates specifically bind to HER2+ cancer cells and have enhanced killing effect due to proximity effects; (**B**) Analysis of HER2 antigen binding on Hereceptin-NK-92MI conjugates. Mean ± SD (error bars); Flow cytometry analysis (**C**) and fluorescent microscopy images (**D**) of specific binding between Herceptin-NK-92MI conjugates and BT474 (HER2+); NK-92MI cells were stained with CellTracker Orange (red), and BT474 cells were stained with CellTracker Green (green). The merged channels of fluorescence and phase contrast are shown; the green fields are clusters of BT474 cells. Scale bar: 50 μm; (**E**) LDH release assay of quantifying cell-mediated cytotoxicity of NK-92MI cells against BT474 cells; Herceptin-NK-92MI conjugates were compared with parental NK-92MI with or without additionally added free Herceptin (5 μg/ml). hIgG-NK-92MI conjugates were used as a negative control. Mean ± SD (error bars), representative graph from three independent experiments; (**F**) Killing activity of Herceptin-NK-92MI conjugates in different GF-Herceptin concentrations for enzymatic transfer. Mean ± SD (error bars); (**G**) Comparison of NK-92MI and Herceptin-NK-92MI conjugates in killing different cancer cell lines with or without HER2 expression. Mean ± SD (error bars), representative graph from three independent experiments; (**H**) Comparison of NK-92MI and Herceptin-NK-92MI conjugates in killing BT474 for different effector to target cell ratios. Mean ± SD (error bars); (**I**) Herceptin and α-EGFR dual labeled NK-92MI cells were compared with Herceptin-NK-92MI conjugates and α-EGFR-NK-92MI conjugates in killing HER2+/EGFR+ SKOV3 cancer cells. Mean ± SD (error bars); (**J**) Comparison of non-irradiated and irradiated (6 Gy) NK-92MI cells in killing BT474 cells; (**K**) *In vivo* anti-tumor activity of Herceptin-NK-92MI conjugates; NSG mice were injected intravenously with 0.5 million MDA-MB-435/HER2+/F-luc cells. Then, the animals were treated once by *i.v.* injection of 3 million NK-92MI or Herceptin-NK-92MI cells on day 1 after the injection of the tumor cells. The control mice received HBSS. Six days after the tumor challenge, the mice were injected with *i.p.* with D-luciferin and imaged by IVIS system. The sizes of the tumors of the mice and mean values ± SD are shown; n = 10. Representative images also are shown. In all figures, ns, P > 0.05; ^*^P < 0.05; ^**^P < 0.01; ^***^P < 0.001; ^****^P < 0.0001.

**Figure 4.**
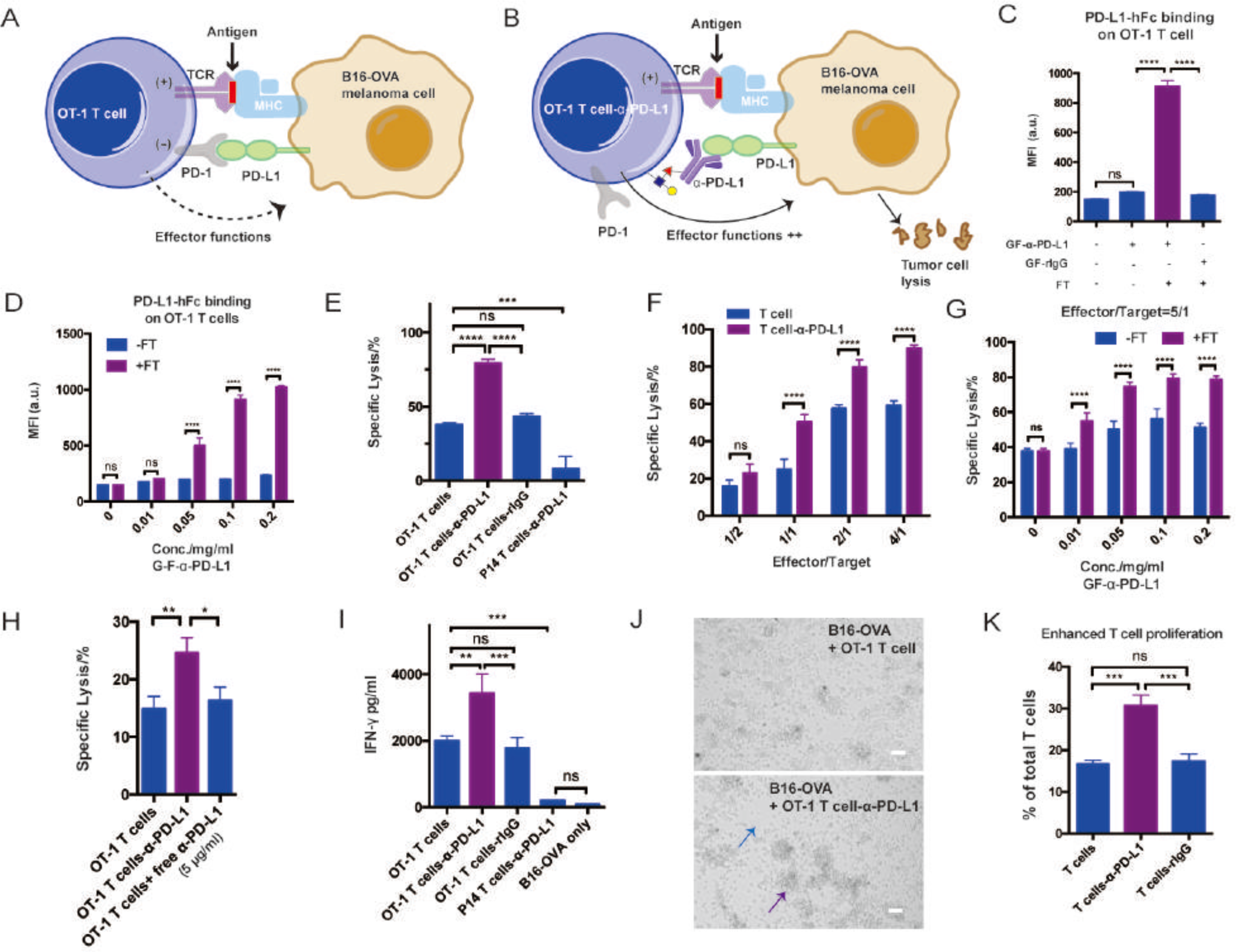
Enzymatic transfer of α-PD-L1 to OT-1 T cells for enhanced T cell activation in specific killing: (**A**) Schematic illustration of the interaction between OT-1 T cells and B16-OVA melanoma cells. MHC complex on B16-OVA present OVA antigen to OVA-specific TCR on OT-1 T cells to induce activation, while PD-L1 on B16-OVA interact with PD-1 on OT-1 T cells to inhibit the activation signal; (**B**) Scheme illustration of the blockade of PD-1/PD-L1 pathway via α-PD-L1 conjugated on the surfaces of OT-1 T cells. The *in situ* blockade could enhance T cell activation and the killing of cancer cells; (**C**) Analysis of PD-L1 antigen binding on OT-1 T cells for different treatments. Mean ± SD (error bars); (**D**) Analysis of PD-L1 antigen binding at different GF-α-PD-L1 concentrations in the enzymatic reaction buffer. Mean ± SD (error bars); (**E**) Assay of quantifying cell-mediated cytotoxicity of α-PD-L1-OT-1 T cell conjugates on B16-OVA cells. Mean ± SD (error bars), representative graph from three independent experiments. OT-1 T cells conjugated with rIgG and P14 T cells conjugated with α-PD-L1 were shown as negative control; (**F**) Comparison of OT-1 T cells and α-PD-L1-OT-1 T cell conjugates in killing B16-OVA at different effector-to-target cell ratios. Mean ± SD (error bars); (**G**) Killing activity of OT-1 T cells and α-PD-L1-OT-1 T cell conjugates for different GF-α-PD-L1 concentrations the enzymatic reaction buffer. Mean ± SD (error bars); (**H**) Comparison of α-PD-L1-OT-1 T cell conjugates with OT-1 T cells with or without additionally added free α-PD-L1 (5 μg/ml) in killing B16-OVA (three hours incubation). (**I**) IFN-γ ELISA of OT-1 T cells mixed with B16-OVA with different treatments. P14 T cell conjugated with α-PD-L1 was shown as negative control. Only B16-OVA was background. Mean ± SD (error bars), representative graph from three independent experiments; (**J**) Microscopy images of OT-1 T cells killing B16-OVA with or without α-PD-L1 labeling. The blue arrow indicates less cancer cells, and the purple arrow indicates larger clusters of T cells. Scale bar: 50 μm; (**K**) Analysis of OT-1 T cell proliferation after the activation mediated by B16-OVA through CFSE dilution. Mean ± SD (error bars);. In all figures, ns, P > 0.05; ^**^P < 0.01; ^***^P < 0.001; ^****^P < 0.0001.

The promising results of enhanced *ex vivo* killing ability of Herceptin-NK92-MI conjugates led us to test their efficacy *in vivo*. Since NK-92 is developed from a patient with lymphoma, as a safety measure, it is usually irradiated prior to clinical use to prevent permanent engraftment in human body. As expected, irradiation of 6 Gy prevented the proliferation of NK-92MI cells (**Supplementary Fig. 16A**). We found gamma irradiated NK-92MI cells maintained their cytotoxicity (**Fig. 3J**) and the half-life of Herceptin conjugated to irradiated NK-92MI was slightly longer than that of the non-irradiated group (**Supplementary Fig. 16B**). After being injected in mice for 24 hr, there was still an abundant amount of Herceptin present on the surface of NK92-MI cells. (**Supplementary Fig. 16C**). To evaluate the *in vivo* efficacy of Herceptin-NK92-MI conjugates, we chose an experimental lung metastasis model in which NSG mice received intravenous (*i.v.*) injections of MDA-MB-435/HER2+/F-luc cells (stably transduced with firefly luciferase). One day after being inoculated with the tumor cells, the mice were treated by *i.v.* injections of irradiated parental NK-92MI cells or Herceptin-NK-92MI conjugates while non-treated group was injected with Hank’s Balanced Salt Solution (HBSS). Six days after tumor inoculations, the volumes of the tumors in the lungs were determined by longitudinal, non-invasive bioluminescence imaging. While treatment with parental NK-92MI cells only moderately reduced the formation of tumors in the lungs, (∼ 48% less than the HBSS group), Herceptin-labeled NK-92MI cells exhibited significantly enhanced *in vivo* tumor killing activity (∼83% less than the HBSS group) (**Fig. 3K**). To assess if Herceptin-NK92-MI conjugates were effective to treat established tumors, NSG mice were injected with luciferase-bearing MDA-MB-435/HER2+ cells intravenously. Three days later, the growth of tumor cells in the lung region was revealed by 3fold increase in bioluminescence (**Supplementary Fig. 17**). The mice were then treated by *i.v.* injections of irradiated parental NK-92MI cells or Herceptin-NK-92MI conjugates on day 3 and day 9. Bioluminescence imaging showed that tumor cell growth was significantly suppressed by Herceptin-labeled NK-92MI cells compared to unmodified NK-92MI cells (**Supplementary Fig. 17**).

### Anti-PD-L1 (α-PD-L1) conjugated on the surfaces of cells could block the PD-1/PD-L1 pathway and enhance the proliferation of T cells *ex vivo*.

Even after infiltrating the tumor bed, cytotoxic functions of effector cells may be dampened by factors produced in the microenvironment of the tumor^27^. As another application of our new technique, we sought to determine whether CD8+ T cells modified by cell-surface mAb conjugation could counteract such inhibitory signals to maintain their activities. The interaction between programmed death 1 (PD-1) receptor, found on T cells, and PD-Ligand (PD-L) expressed by tumor cells plays a major role in inhibiting the cytotoxicity of T cells^28^ (**Fig. 4A**). We hypothesize that the installation of α-PD-L1 on the surfaces of T cells may block the PD-1/PD-L1 interaction *in situ* to enhance the activation of the T cells and thus enforcing tumor cell lysis (**Fig. 4B**).

OT-1 transgenic mice was chosen as the model system whose CD8+ T cells express a T cell receptor (TCR) specific for the SIINFEKL peptide (OVA_257-264_) of ovalbumin presented on MHC I. Splenocytes from the OT-1 mice were first stimulated with OVA_257-264_ peptide and expanded *ex vivo* in the presence of IL2 or IL7/IL15. After three days, most of the cells are activated OT-1 CD8+ T cells (**Supplementary Fig. 18A**). During the next 10 days, OT-1 T cells were confirmed to express high levels of LacNAc (**Supplementary Fig. 18B**) and were subjected to the chemoenzymatic modification with GF-rIgG (**Supplementary Fig. 18C**). Re-stimulation experiments confirmed that the modified T cells had a similar proliferation rate to that of the unmodified cells, suggesting that the conjugated IgG molecules do not block the interaction between TCR and the MHC I-complex (**Supplementary Fig. 18D**).

Subsequently, CD8+ T cells from OT-1 or P14 mice, which carry a transgenic TCR that recognizes the gp33-41 epitope of lymphocytic choriomeningitis virus, were modified with GF-α-PD-L1 using chemoenzymatic glycan engineering (**Supplementary Figs. 19A and 19B**). α-PD-L1 maintained its antigen-binding capacity upon cell-surface conjugation (**Fig. 4C and Supplementary Fig. 19C**). As expected, the degree of both cell-surface conjugation and antigen binding were dependent on the concentration of GF-α-PD-L1 (**Fig. 4D and Supplementary Fig. 19D**). At the saturated condition, approximately 7×10^4^ anti-PD-L1 molecules were conjugated to the cell surface (**Supplementary Fig. S6D**). Then, the modified OT-1 CD8+ T cells were subjected to an *ex vivo* killing assay, in which a B6-derived melanoma cell line B16F10 expressed ovalbumin (B16-OVA) was used as model target cells. After incubation for 20 hours, the OT-1 T cells conjugated with α-PD-L1 showed significantly enhanced lysis of B16-OVA cells (**Fig. 4E**), and the enhanced killing effect was observed only when the effector-to-target cell ratio was above 1 (**Fig. 4F**). By contrast, the specific lysis of B16-OVA induced by the rIgG-OT-1 T cell conjugates was much weaker, which is similar to that of the control, unmodified T cells (**Fig. 4E**). CD8+ T cells of irrelevant specificity from P14 mice also were conjugated with α-PD-L1 as a negative control. Remarkably, this ACC only induced background killing (**Fig. 4E**), suggesting that trace levels of α-PD-L1 conjugated to the surfaces of the cells could not mediate significant target cell lysis. Although the antigen binding capacity reached the maximum when 0.1 mg/ml GF-α-PD-L1 was used for cell-surface conjugation (**Fig. 4D**), the optimal killing capacity was achieved at ∼ 0.05 mg/ml GF-α-PD-L1 (**Fig. 4G**), suggesting that only half of the maximum cell-surface conjugation is required for efficient blocking the PD-1–PD-L1 interaction. Moreover, using the same killing assay we observed that α-PD-L1-modified OT-1 cells exhibited significantly better lysis capability than a simple combination of OT-1 cells with free α-PD-L1 (**Fig.4H**).

To further determine whether the cell-surface conjugated α-PD-L1 could suppress PD-1/PD-L1 co-inhibitory signaling, we measured the cytokine produced by modified OT-1 T cells when they were mixed with B16-OVA. The enhanced IFN-γ and TNF-α secretion only was observed in T cells conjugated with α-PD-L1, which exhibited dependency on the dosage of conjugated α-PD-L1 (**Fig. 4I and Supplementary Fig. 20**). The enhanced T-cell activation also was observed directly using microscopy due to the formation of larger clusters of T cells in the α-PD-L1 labeled group (**Fig. 4J and Supplementary Fig. 21A**). Finally, the enhanced activation of T cells also promoted T cell proliferations as confirmed by a CFSE dilution assay by re-stimulating modified OT-1 T cells with B16-OVA (**Fig. 4K and Supplementary Fig. 21B**).

## Conclusion

The one-step FucT-based chemoenzymatic method developed here is a fast, simple, and cost-effective technique for cell engineering, especially for constructing ACCs. Using this method, we successfully constructed two ACCs, which exhibited enhanced activities in two critical stages of anti-cancer immune responses, *i.e.*, targeting and killing. Although mAbs conjugated on the cell surface are diluted due to internalization and cell proliferation (cell-surface half-life ∼ 8-24 h in our experiments), this method offers several advantages over genetic-based engineering, *e.g.* rapid and homogeneous modification, capability to install multiple mAbs simultaneously, and therefore serves as a nice complement or synergistic method to the permanent, genetic engineering approach that has found great success in making CAR constructs, including CAR-NKs based on NK-92 cells^29^. In these cases, γ-irradiation is employed as a potential safety measure for clinical application to prevent NK cell replication while preserving their antitumor activities^29^. Upon irradiation, NK cells can no longer replicate. Therefore, there is no obvious advantage of using these CAR constructs than using ACCs disclosed here. The fact that NK-92MI cells are currently undergoing clinical trials and Herceptin is already a FDA-approved drug heralds the potential of Herceptin-NK-92MI conjugates for further development as a clinical candidate. More broadly, our method is easily generalized to conjugate NK-92MI cells with other therapeutic mAbs to achieve synergistic effects for different kinds of target in cancer immunotherapy.

## Acknowledgments

This work was supported by the NIH (GM113046 and GM093282 to P.W.). We thank Prof. Carolyn Bertozzi (Stanford, USA) for NK-92MI cells, Prof. Peter Schultz (TSRI, USA) for MDA-MB-435/HER2+/F-luc cells, Prof. Peng R. Chen (PKU, China) for the Sortase A plasmid, Prof. John Teijaro (TSRI, USA) for P14 mice and Prof. Philippe A. Gallay (TSRI, USA) for NSG mice.

## Author contributions

P.W., J.L., and M.C. designed the experimental strategy and wrote the manuscript. J.L. and M.C. performed the experiments with the help of L.Z., Z.L., and M. A. P.W. and J.L. prepared the figures, and all authors edited the manuscript.

## Additional information

Supplementary information and chemical compound information is available online. Correspondence and requests for materials should be addressed to P.W.

## Supplementary Information

## General Considerations

### Materials and Reagents

All chemical reagents and solvents were obtained from Sigma-Aldrich and used without further purification unless otherwise noted. All cell culture materials are listed in cell culture methods. ImmunoCult^TM^ human CD3/CD28 T cell activator was purchased from STEMCELL Technologies, Inc. Recombinant mouse IFN-γ, recombinant mouse PD-L1-human Fc Chimera (PD-L1-hFc), CFSE cell division tracker kit, APC Streptavidin, anti-mouse CD3 (17A2)-FITC, anti-mouse CD8 (53-6.7)-PE, anti-mouse CD45.1 (A20)-Pacific blue, anti-mouse Thy1.1 (OX7)-Alexa Fluor 700, anti-mouse IgG (Poly4060)-APC, anti-human Fc (HP6017)-APC, anti-rat IgG (Poly4050)-APC, anti-His Tag (J095G46)-PE, anti-human CD3 (HIT3a)-APC, anti-human CD45 (HI30)-FITC, anti-human CD4 (RPA-T4)-PE/Cy7, anti-human CD8 (SK1)-Pacific Blue, anti-human CD25 (M-A251)-PerCP/Cy5.5, anti-human CD44 (BJ18)-FITC, anti-human CD45RO (UCHL1)-Alexa Fluor 700, anti-human CD62L (DREG-56)-PE and human Fc Receptor blocking solution were purchased from Biolegend. Bulky monoclonal antibodies including isotype mouse IgG2a (C1.18.4), anti-mouse PD-L1 (α-PD-L1, 10F.9G2), isotype rat IgG2b (LTF-2) and anti-human EGFR (528) were purchased from Bio X Cell. Therapeutic Herceptin were from Genentech. The control human IgG was obtained from Athens Research and Technology. Recombinant human HER2/ErbB2 Protein with His Tag (HER2-His) was purchased from Sino Biological, Inc. Click reagents including biotin-PEG3-azide (AZ-104, MW: 444.5), methyltetrazine-PEG4-azide (1014, MW: 389.40), methyltetrazine-PEG4-alkyne (1013-old, MW: 487.5, a discontinued product in their website), TCO-PEG4-NHS Ester (A137, MW: 514.6) and Cy5-TCO (1089, MW: 959.20) were purchased from Click Chemistry Tools LLC. Cy3-Azide (MW: 712.8) is a gift from Prof. Xing Chen’s lab (PKU, China). (5OctdU)-5’-CAGTCAGTCAGTCAGTCAGT-3’(6-FAM) was ordered from Integrated DNA Technologies, Inc. D-Luciferin (monosodium salt), TNF-α mouse ELISA kit, IFN-γ mouse ELISA kit, Granzyme B human ELISA kit, DiD’ solid, Hoechst 33342, DAPI, Alexa Fluor 647 NHS Ester, cell tracker green (CMFDA) and orange (CM^TM^R) were purchased from Thermo Fisher Scientific. 5×RIPA buffer kit were purchased from Cell Biolabs, Inc. CytoTox 96® non-radioactive cytotoxicity assay and Bright-Glo^TM^ luciferase assay system were purchased from Promega Corporation. Inorganic pyrophosphatase, FKP and FucT were expressed in endotoxin free bacteria (ClearColi® BL21-DE3, Lucigen) and purified as previously reported^1^. Pure GDP-Fuc, GF-Al and BTTP were synthesized as previously described ^1,2^. The plasmid of Sortase A (5M) is a kind gift from Prof. Peng Chen (PKU). The enzyme was expressed and purified according to their work ^3^.

### Equipment

All of the flow cytometry analyses were performed on an Attune NxT Flow Cytometer. Images of protein gels including coomassie SDS-PAGE gel and western blotting membrane were taken on ChemiDoc XRS+ (Bio-Rad). Absorbance, fluorescence intensity and luminescence were monitored in a Multi-Mode Microplate Reader (Synergy^TM^ H4, Bio-Tek). Confocal images were taken on a Nikon spinning disk confocal microscope (TE2000). Fluorescent and phase contrast microscope images were taken on an All-In One fluorescence microscope (Keyence, BZ-X700). Bioluminescence imaging of live mice were acquired using an IVIS Spectrum system.

### Cells

Cell lines were all purchased from ATCC unless otherwise specified. CHO cell lines (WT, Lec2 and Lec8, from Prof. Pamela Stanley at Albert Einstein College of Medicine) were grown as monolayer in alpha-Minimum Essential medium (α-MEM) (GIBCO) supplemented with 10% fetal bovine serum (FBS) (Omega Scientific, Inc). NK-92MI cell line was grown in MyeloCult^TM^ H5100 (STEMCELL Technologies, Inc). Cancer cell lines including BT474, SKBR3, MDA-MB-435 (HER2+ and HER2-), MDA-MB-468, SKOV3 and mouse B16-OVA (from Prof. Gregoire Lauvau lab) are all grown in grown in DMEM (Dulbecco’s modified Eagle’s medium, GlutaMAX, GIBCO) supplemented with 10% FBS. Human blood samples were collected from healthy donors under the TSRI Normal Blood Donor Services program (#IRB 15-6710). Peripheral blood mononuclear cells (PBMCs) were obtained by Ficoll (Ficoll-Paque Plus, GE) density centrifugation. PBMC, activated human T cells and mouse T cells were all cultured in RPMI 1640 (GlutaMAX) with 10% heat-inactivated FBS, 1 mM sodium pyruvate, 50 μM β-ME, 10mM HEPES and 1×MEM NEAA (GIBCO) (referred as T cell culture media later). Cytokines in T cell culture media were added as indicated (rhIL2, rhIL7 and rhIL15 are all from NIH program). All cells cultures were incubated at 37 °C under 5% CO_2_.

### Mice

All mice were bred or housed under specific pathogen free (SPF) conditions. All animal experiments were approved by TSRI Animal Care and Use Committee. OT-1 mice are purchased from Taconic Biosciences. CD 45.1 mice from C57BL/6J (B6) genetic background were purchased from the Jackson Laboratory. The strain of OT-1+/-/CD 45.1+/-was generated by cross breeding. Six female B6 background P14 mice were gifts from Prof. John Teijaro lab. 176 female NSG mice are gifts from Prof. Philippe A. Gallay lab. Both male and female mice of 8–20 weeks of age were used for most experiments.

## Supplementary Methods

### 1. One-pot protocol for producing GDP-Fuc derivatives

Reactions were typically carried out in a 15 mL corning tube with 5 mL 100 mM HEPES buffer (pH 7.5) containing L-fucose analogues (final concentration, 10 mM), ATP (10 mM), GTP (10 mM), MgSO_4_ (10 mM), KCl (50 mM), inorganic pyrophosphatase (90 units, ∼0.17 g/L, endotoxin free), and FKP (9 units, ∼0.6 g/L, endotoxin free). The reaction mixture was incubated at 37 °C for 5–6 h with shaking (225 rpm). After the reaction finished (monitored by TLC analysis^1^), enzymes were precipitated by adding 5 mL cold EtOH into the crude product. After the precipitates were removed through centrifuge (8000×g, 5 min), the crude products (containing ∼10 mM GDP-Fuc analogues) could be directly used. For GF-Al and GF-Az, further modification could be achieved through CuACC reaction. Crude GF-Al/GF-Az sample (∼5 mM, in HEPES buffer) were reacted with azide/alkyne probes (6 mM) in the presence of Cu/BTTP (1/2, 500 μM) and sodium ascorbate (2 mM) at 30 °C for 6h (For Tz substrates, add one volume of MeOH in reaction mixture). After reaction finished (monitored by TLC and LC-MS analysis, representative figures of GF-Az-Tz and GF-Al-Tz are shown in **Spectra** part), BCS (bathocuproine sulphonate, 2 mM) were added to quench the reaction. These biocompatible crude products were lyophilized and reconstituted in pure water to a final concentration of 10 mM, which could be directly used in cell labeling. Following this protocol, we made one-pot products of GF-Al-Biotin, GF-Al-Cy3, GF-Al-Tz and GF-Az-Tz. Their structures are shown in ***Figure S1***. Since GF-Al-Tz and GF-Az-Tz are important starting materials for making GF-IgG, we purified these two GDP-fucose derivatives and characterize them using high-resolution ESI-TOF MS and NMR (**Spectra**), which also confirm the one-pot procedure is reliable.

### 2. General procedure for enzymatic transfer of GDP-Fuc derivatives to cell surface

Live cells (1∼2 million) were resuspended in 100 μL HBSS buffer containing 20 mM MgSO_4_, 3 mM HEPES, 0.5% FBS, 100 μM GDP-Fuc derivatives and 30 mU FucT (∼0.02 mg/mL). After the incubation for 20 minutes (works from on ice to 37 °C), the cells were washed with PBS and ready for further application or analysis. For biotin detection, the cells were stained with APC Streptavidin after reaction. For Tz detection, the cells were reacted with 20 μM TCO-Cy5 on ice for 30 minutes and washed three times.

### 3. Comparison of Sortagging and Fucosylation

For sortagging reaction, NK-92MI cells (5 million/ml) were incubated with soluble SrtA and biotin-LPETG in RPMI complete media (10% inactivated fetal bovine serum, 1% MEM (Gibco) Non-Essential Amino Acids Solution, 1 mM sodium pyruvate, 1% GlutaMAX, (Gibco) 0.1 mM β-mercaptoethanol) supplemented with 10 mM CaCl_2_ at 37 °C for 2 hr^4^. For fucosylation reaction, NK-92MI cells (5 million/ml) were incubated with FucT and GF-biotin in fucosylation buffer described in Methods 2 for different times (2-30 min). Different concentrations of enzymes and substrates were used as indicated.

### 4. Preparation of GDP-Fuc modified antibodies

All of the antibodies (full-length IgG, MW: ∼150 KDa) for conjugation were first desalted into PBS and concentrated to a 6 mg/mL solution. TCO group was first introduced onto antibodies according to the standard labeling protocol of TCO-PEG4-NHS ester (https://clickchemistrytools.com/product/tco-peg4-nhs-ester/) and previous reports^5,6^ about its application on IgG labeling. Briefly, we prepared fresh 50 mM stock of TCO-PEG4-NHS reagent in DMSO and add it to the IgG sample (final concentration 6 mg/mL) at a final concentration of 0.5 mM. The reactions were incubated at room temperature for 30 minutes and quenched by adding Tris buffer (pH 8.0) to a final concentration of 50mM Tris. The quenched reaction mixtures were incubated at room temperature for 5 minutes and then desalted into PBS using G25 desalting column (PD-10, GE). The concentrations of desalted TCO-IgGs were around 4.5 mg/mL (∼1-4 TCO per IgG). After that, ∼10 mM one-pot products of GF-Az-Tz were added to TCO-IgGs with a final concentration at 0.15 mM (5eq of IgG). After 30 minutes of incubation at room temperature, these one-pot GF-IgG products were ready to use and could be kept at 4 °C for up to 2 months. All the GDP-fucose modified antibodies were characterized by MALDI-TOF MS. The molecular weights and average numbers of GF modification were shown in **Supplementary Figure S5B**. To make Alexa Fluor 647 labeled GF-IgGs used in confocal imaging and fluorescent gel analysis, the GF-IgG molecule was labeled with Alexa Fluor 647 probes following the manual. The antigen binding affinities of GF-Herceptin, GF-anti-EGFR and GF-anti-PD-L1 were titrated on MDA-MB-435 Her2, SKOV3 and B16-OVA (IFN-γ treated) cells, respectively, according to the reference ^5^.

### 5. General procedure for enzymatic transfer of GF-IgG to cell surface

Live cells (1∼2 million) were resuspended in 100 μL HBSS buffer containing 20 mM MgSO_4_, 3 mM HEPES, 0.5% FBS, 0.1 mg/mL one-pot GF-IgG (different concentrations were used in titration) and 60 mU FucT (∼0.04 mg/mL). After the incubation for 20 minutes (works from on ice to 37 °C), the cells were washed with PBS and ready for further application or analysis. For labeling detection, IgG labeled cells were stained with DAPI and fluorescent secondary antibody against labeled IgG (1/50-1/200 dilution) on ice for 30 minutes. For confocal imaging, Lec2 cells plated on a chamber cover glass (Nunc) were treated with 0.1 mg/mL Alexa Fluor 647 labeled GF-rIgG in the same fucosylation condition as described above. After conjugation, live cells were washed and stained with Hoechst 33342 for 30 minutes on ice and then washed for imaging. To confirm the binding activity of cell surface conjugated antibodies, labeled cells were allowed to bind with 10 μg/mL antigen (hE-sel-hFc, PD-L1-hFc or HER2-His) in binding buffer (HBSS with 5mM HEPES, 2mM CaCl_2_, and 1mM MgCl_2_) on ice for 30 minutes. After binding, cells were washed twice with PBS and stained with DAPI and APC-anti-human Fc (1/50 dilution) or PE-anti His (1/100 dilution). For the SDS-PAGE fluorescent gel imaging analysis, GF-IgG-A647 was used instead of GF-IgG. At the saturated labeling condition, labeled cells and control cells were collected and lysed in SDS loading buffer and subjected to SDS-PAGE analysis. Pure GF-IgG-A647 proteins were also loaded as standards in quantification.

### 6. Primary human T cells preparation and IgG labeling on their surface

Fresh PBMCs were freshly prepared as described above. 4 million per mL PBMCs were cultured in T cell culture media with 15 ng/mL rhIL2 and activated with human CD3/CD28 T cell activator for two days. After that, activated human T cells were kept under 4*10^6^ cells/mL in T cell culture media (fresh media with cytokine were added every two days). Phenotypes were characterized after two weeks expansion (>95% are human T cells). LacNAc levels on CD4+ and CD8+ T cells were tracked through fucosylation with GF-biotin on day 0 (naïve T cells), day 2, day 4, day 7, day 11 and day 13. Activated human T cells were labeled with α-hE-Sel and mIgG control using the general procedure for enzymatic GF-IgG transfer. After that, the labeling detection and the antigen (hE-sel-hFc) binding were both confirmed. Labeled human T cells were then cultured in T cell culture media at a start cell density of 0.5*10^6^ cells/mL. The decay of cell surface mIgG molecule was tracked in 24 hours after labeling (anti-mouse IgG staining). The cell proliferation rate of labeled T cells was compared with unlabeled human T cells in three days (live cell counting).

### 7. IgG labeling on NK-92MI cells

NK-92MI cells or irradiated NK-92MI cells (6 Gy) were labeled with Herceptin or control human IgG according to the general protocol. After labeling, the labeling detection and the antigen (HER2-His) binding were both confirmed. NK-92MI cells with or without IgG conjugation were then cultured in T cell media with a start cell density of 0.5*10^6^ cells/mL. The decay of ell surface conjugated Herceptin were tracked at day 0, day 1 and day 2 (anti-human Fc staining). The proliferation rates of cells were tracked at day 0, day 1, day 2 and day 3 (DAPI staining and FACS counting). For dual antibodies labeling, NK-92MI cells were conjugated with GF-α-EGFR first and then conjugated with GF-Herceptin after washing.

### 8. FACS and imaging analysis of binding between NK-92MI and BT474

For flow cytometry analysis, NK-92MI cells were stained with CFSE and then labeled with Herceptin or not according to the general protocol. BT474 cells were stained with DiD first and then mixed with Herceptin labeled or unlabeled NK-92MI cells at the ratio of 1:1. Two hours later, the cells mixture was analyzed by flow cytometry. For fluorescence imaging, BT474 cells were stained with CFSE and cultured in glass bottom petri dish overnight. NK-92MI cells were then stained with cell tracker orange and then labeled with Herceptin or not. NK-92MI cells were added to BT474 culture at ratio of 1:1. Two hours later, the co-cultured cells were imaged by fluorescent microscope before and after PBS wash.

### 9. Analysis of NK-92MI cells-mediated cytotoxicity against HER2+ cancer cells

Labeled or unlabeled NK-92MI cells were co-cultured with different type of cancer cells at indicated effector/target ratios for 4 hours in a 96-well plate. In most of the experiments, the effector/target ratio is 5/1. Free Herceptin were added at a final concentration of 5 μg/mL if indicated. Specific cancer cell lysis was detected by LDH secretion in supernatant (CytoTox 96, Promega). Set-up of control groups and calculations of specific lysis were according to manufactory’s instruction (https://www.promega.com/products/cell-health-assays/cell-viability-and-cytotoxicity-assays/cytotox-96-non_radioactive-cytotoxicity-assay/?catNum=G1780). Supernatant of each group (one hour incubation) were also collected and subjected to granzyme B ELISA kit for quantification.

### 10. Mice model of NK-92MI mediated killing of HER2+ cancer

Thirty female NSG mice (6-8 weeks old) were inoculated with 5*10^5^ MDA-MB-435 HER2+/F-luc cells through tail vein injection. One day later, mice were randomly divided into three groups (10 mice per group). Each group were treated with HBSS, NK-92MI or Herceptin labeled NK-92MI cells through tail vein injection (3*10^6^ NK cells each mice). Six days after tumor challenge, mice were injected with 200 μL D-luciferin (15mg/mL) through *i.p.* injection. Twelve minutes later, the bioluminescence signal in mice were analyzed by PerkinElmer IVIS system. The total photons indicating the tumor mice were quantified by IVIS software. For the established tumor model, NSG mice were injected intravenously with 5*10^5^ MDA-MB-435/HER2+/F-luc cells on day 0. Three days later, the animals were imaged to confirm the tumor formation and then treated by *i.v.* injection of 5*10^6^ NK-92MI or Herceptin-NK-92MI cells or HBSS. The animals were imaged again on day 4 and day 10. The animals were received one more treatment of 5*10^6^ NK-92MI or Herceptin-NK-92MI cells (*i.v.* injection) on day 9.

### 11. Primary OT-1 CD8+ T cells preparation and IgG labeling on their surface

Splenocytes from OT-1 mice were activated by 1nM OVA peptides in T cell media for two days. After that, activated cells were *in vitro* expanded in fresh T cell media with 15 ng/mL rhIL2 or 10 ng/mL rhIL7 and 20 ng/mL rhIL15 for several days (kept under 8*10^6^ cells/mL, fresh media with cytokine were added every two days). Phenotypes were characterized (>95% are OT-1 CD8+ T cells). LacNAc levels on OT-1 T cells (cultured with two different cytokines) were tracked through fucosylation with GF-biotin on day 0 (naïve T cells), day 2, day 4, day 7, day 9, day 11 and day 13. Activated OT-1 T cells were labeled with α-PD-L1 and mIgG control using the general procedure for enzymatic GF-IgG transfer. After that, the labeling detection and the antigen (PD-L1-hFc) binding were both confirmed. Labeled OT-1 T cells were then cultured in T cell culture media at a start cell density of 0.5*10^6^ cells/mL. The decay of cell surface α-PD-L1 molecule was tracked in 24 hours after labeling (anti-rat IgG staining).

### 12. OT-1 T cells re-stimulation using OVA-pulsed splenocytes

Splenocytes from B6 WT mice were pulsed with SIINFEKL (OVA) peptide (1 μg/mL, in T cell media) for 2h. After that, cells were washed three times before use. OT-1 T cells with CD45.1 congenic marker were stained with CFSE and then labeled with rIgG. 10^5^ OT-1 T cells (labeled or unlabeled) were mixed with 10^6^ OVA-pulsed splenocytes in 500 μL T cell media. Control groups were also set up without splenocytes. The cell mixtures were cultured for 3 days. After that, cells were stained with APC anti-CD45.1 and DAPI. The CFSE dilution signals were analyzed on live CD45.1 positive cells.

### 13. Analysis of OT-1 CD8+ T cells mediated cytotoxicity against B16-OVA

B16-OVA cells (stably transduced with firefly luciferase) were seeded in 96-well plate and treated with 10 ng/mL IFN-γ overnight (B16-OVA cells were all treated with IFN-γ to induce high expression level of PD-1 in this work). Labeled or unlabeled OT-1 cells were co-cultured with B16-OVA cancer cells at indicated effector/target ratios for 3 hours or 20 hours in a 96-well plate (free anti-PD-L1 was added at 5 μg/ml when indicated). In most of the experiments, the effector/target ratio is 5/1. Phenotype of T cell clusters was imaged before cell number quantification. B16-OVA cell numbers were quantified through the luciferase activity according to the reference. The detection reagent was directly added to medium in each well according to the manufactory’s manual (Bright-Glo, Promega). For cytokine secretion quantification, cells were cultured for 9 hours and supernatant were collected and subjected to TNF-α and IFN-γ ELISA kit. For OT-1 T cells proliferation in killing B16-OVA, OT-1 cells were stained by CFSE before IgG labeling. Modified cells were mixed with B16-OVA cells in the effector/target ratio of 2/1. After 72 hours, the cell mixtures were stained with APC-anti CD8 and DAPI. CFSE dilution signal was analyzed on CD8+ cells.

**Figure S1.**
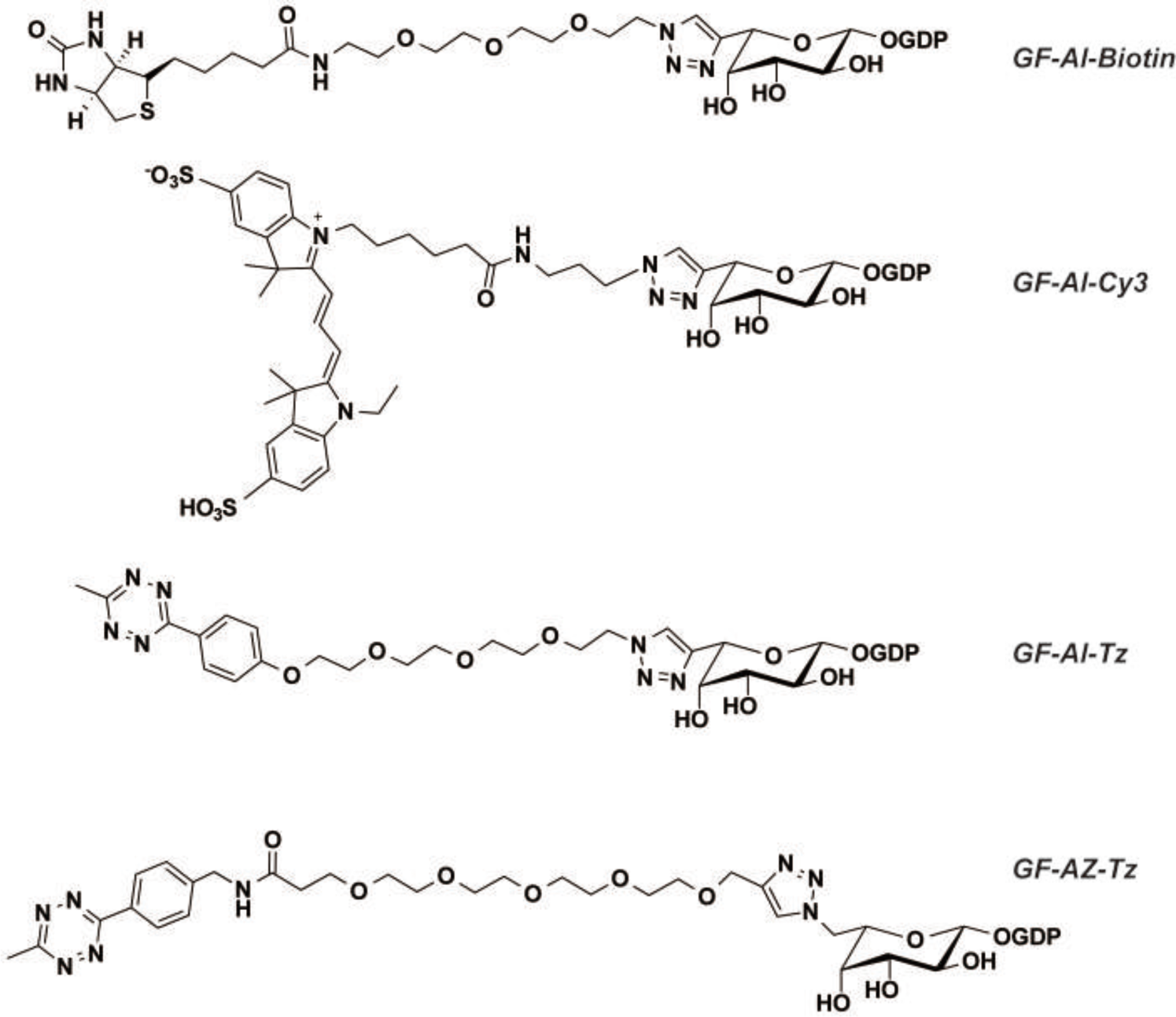
Chemical s*tructure*s of GDP-Fuc derivatives including GF-Al-Biotin, GF-Al-Cy3, GF-Al-Tz and GF-Az-Tz.

**Figure S2.**
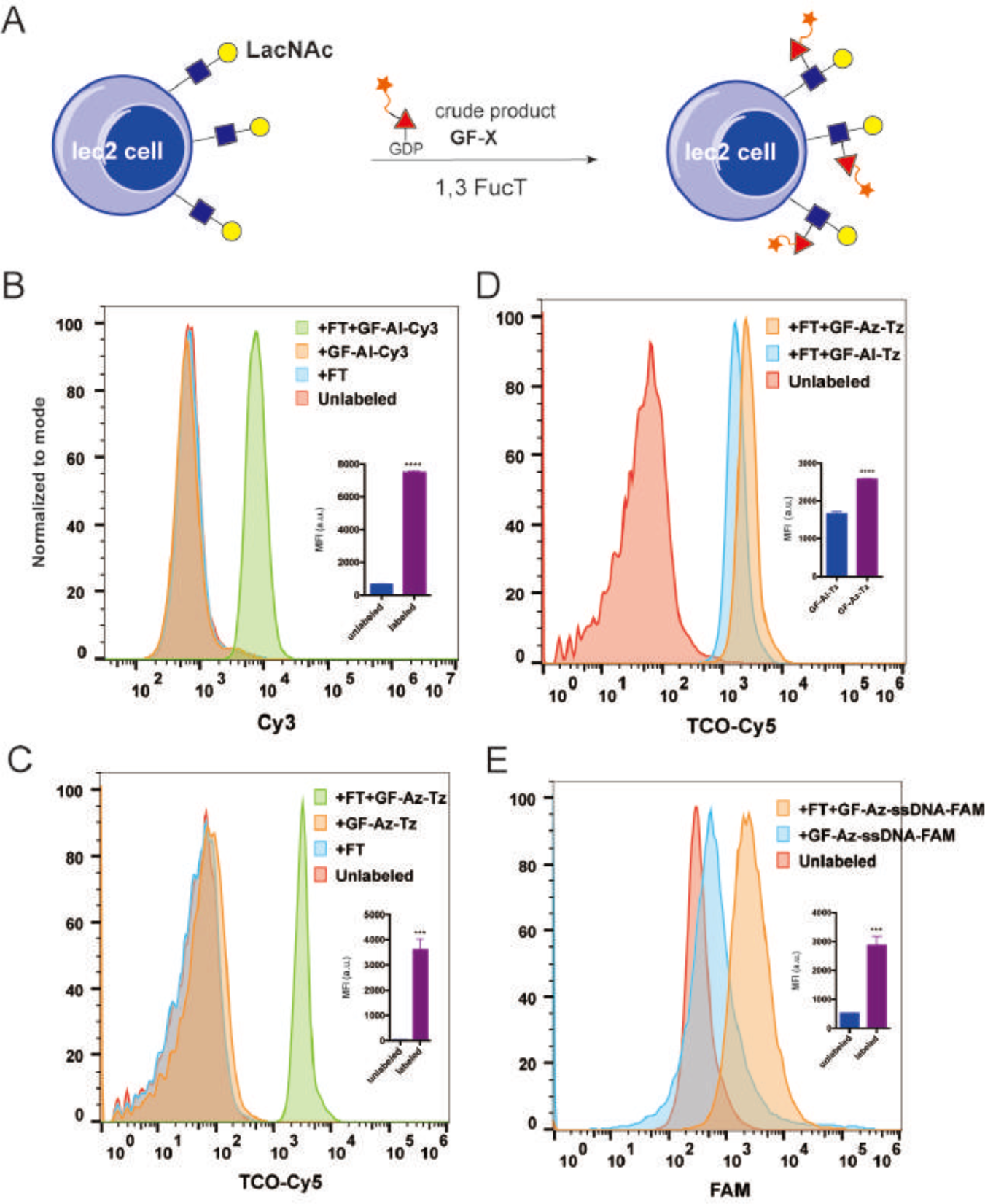
The one-step enzymatic transfer of one-pot GDP-Fu*c* derivatives to Lec2 cells. (**A**) Schematic representation of transferring one-pot GDP-Fuc derivatives using α1,3-FucT to LacNAc epitope on Lec2 CHO cell surface. (**B**) Lec2 cells were treated with FT and GF-Al-Cy3, FT alone, or GF-Al-Cy3 alone, or untreated. Then cells were analyzed by flow cytometry, which only showed successful labeling when treated with both FT and GF-Al-Cy3. (**C**) Lec2 cells were treated with FT and GF-Az-Tz, FT alone, or GF-Az-Tz alone, or untreated. Then the cells were reacted with TCO-Cy5 and analyzed by flow cytometry. Clear labeling is only shown in the group treated with FT and GF-Az-Tz. (**D**) GF-Al-Tz and GF-Az-Tz were compared in the FT mediated transfer reaction on Lec2 cells. Cells were reacted with TCO-Cy5 after enzymatic reaction and analyzed by flow cytometry. GF-Az-Tz group showed stronger signal than GF-Al-Tz. (**E**) Lec2 cells were treated with FT and GF-Az-ssDNA-FAM, or GF-Az-ssDNA-FAM alone, or untreated. Then the cells were analyzed by flow cytometry. The group treated with FT and GF-Az-*ss*DNA-FAM has the strongest FAM signal.

**Figure S3.**
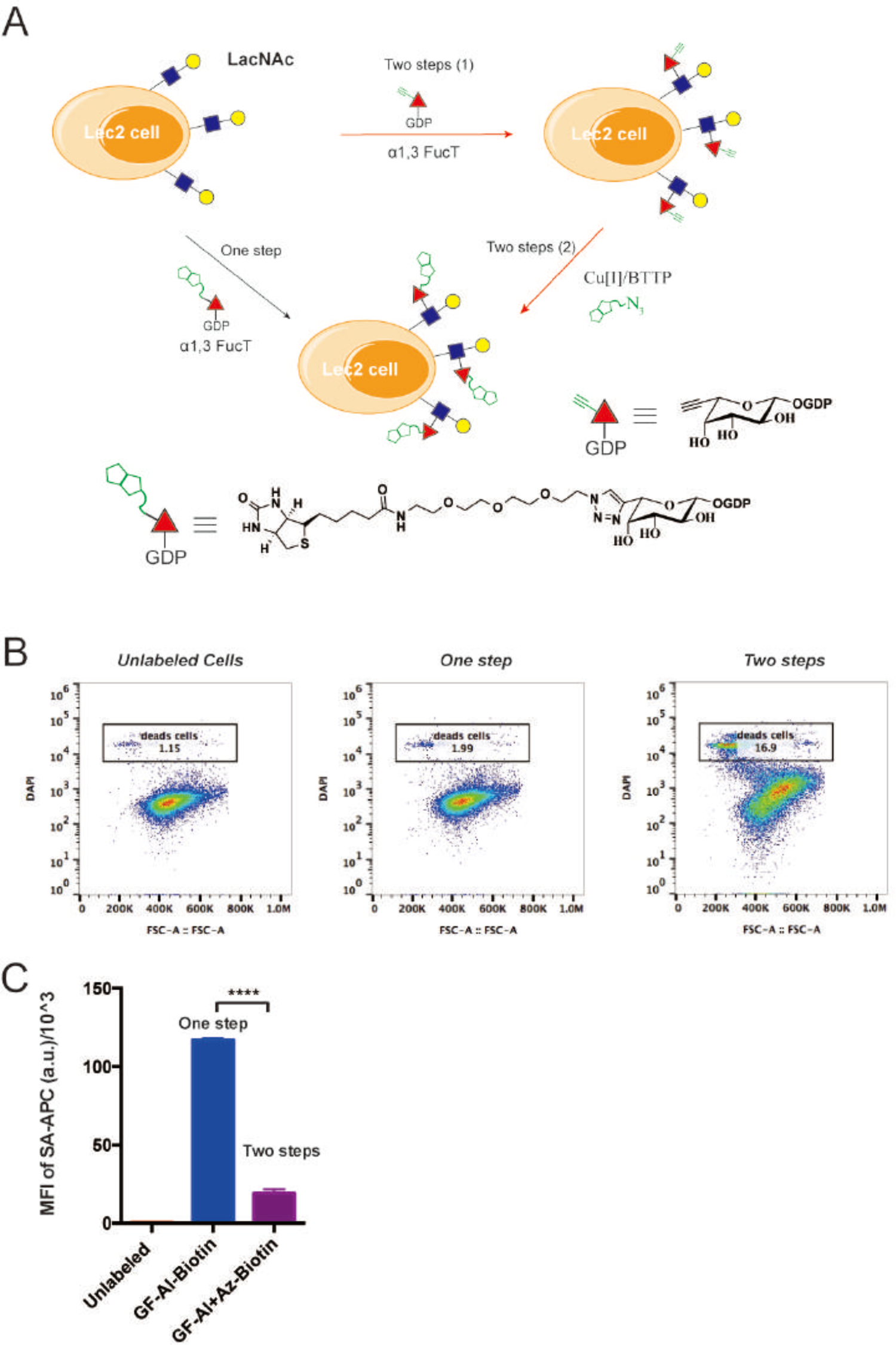
The one-step enzymatic labeling is more efficient and biocompatible than the two-step approach. (**A**) Schematic representation of the one-step and the two-step labeling system. In one step labeling system, one-pot product of GF-Al-biotin was directly transferred to LacNAc epitope on Lec2 cell surface using α1,3-FucT. In contrast, the two-step labeling involved the first step of enzymatic transfer of GF-Al and a followed step of CuAAC reaction between surface alkyne and azide-biotin probe. (**B**) Flow cytometry analysis of biocompatibility in one-step or two-step labeling process. Lec2 cells after reaction were stained with DAPI. (**C**) Comparison of efficiency in one-step and two-step labeling process described in (A). Cells were stained by streptavidin-APC.

**Figure S4.**
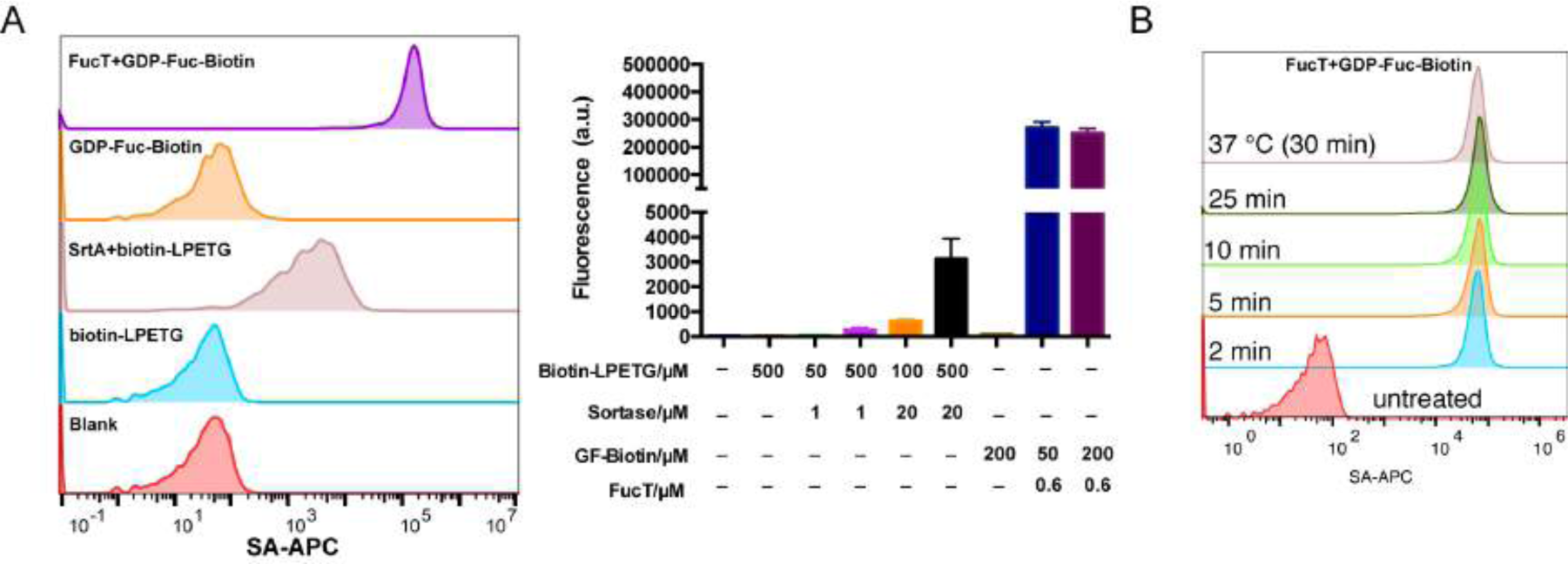
Direct comparison of FucT and Sortase A (5M) for NK-92MI cell surface engineering. (**A**) FT and GF-biotin were used in cell-surface fucosylation and SrtA 5M and biotin-LPETG were used in sortagging. Enzyme alone, biotin probe alone or untreated were shown as negative controls. Different enzyme and probe concentrations were used in the direct comparison experiment. (**B**) Time course of enzymatic transfer of GF-biotin to NK-92 MI cells on ice; reaction at 37 ^º^C was used as the maximum labeling control.

**Figure S5.**
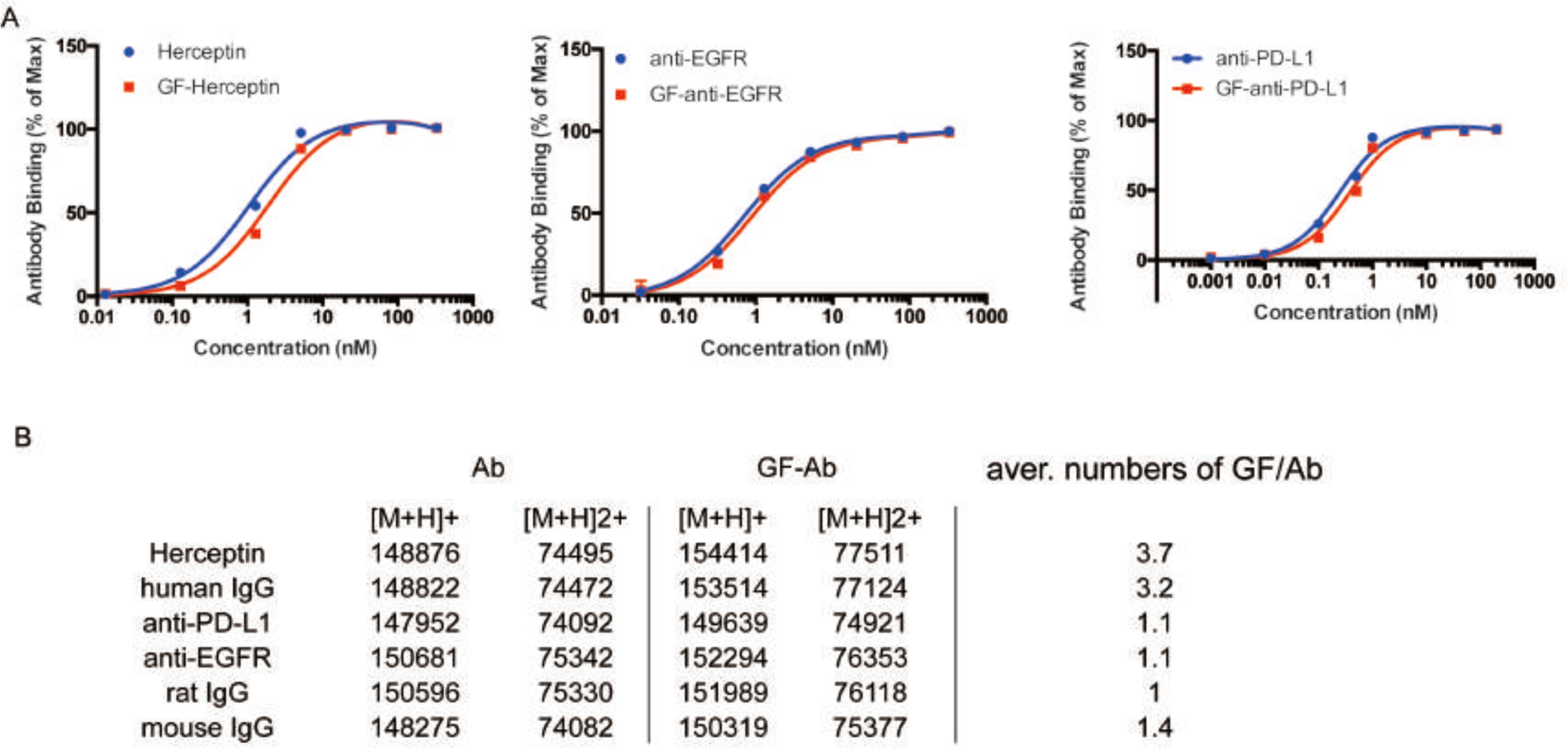
Characterization of GDP-fucose modified antibodies. (**A**) Antigen binding capacities of GDP-fucose modified antibodies were titrated and compared with their parent antibodies. (**B**) Molecular weights of GDP-fucose modified antibodies were characterized by MALDI-TOF MS. Average numbers of GDP-fucose on each antibody were calculated (one GF attachment leads to a 1489 Da shift).

**Figure S6.**
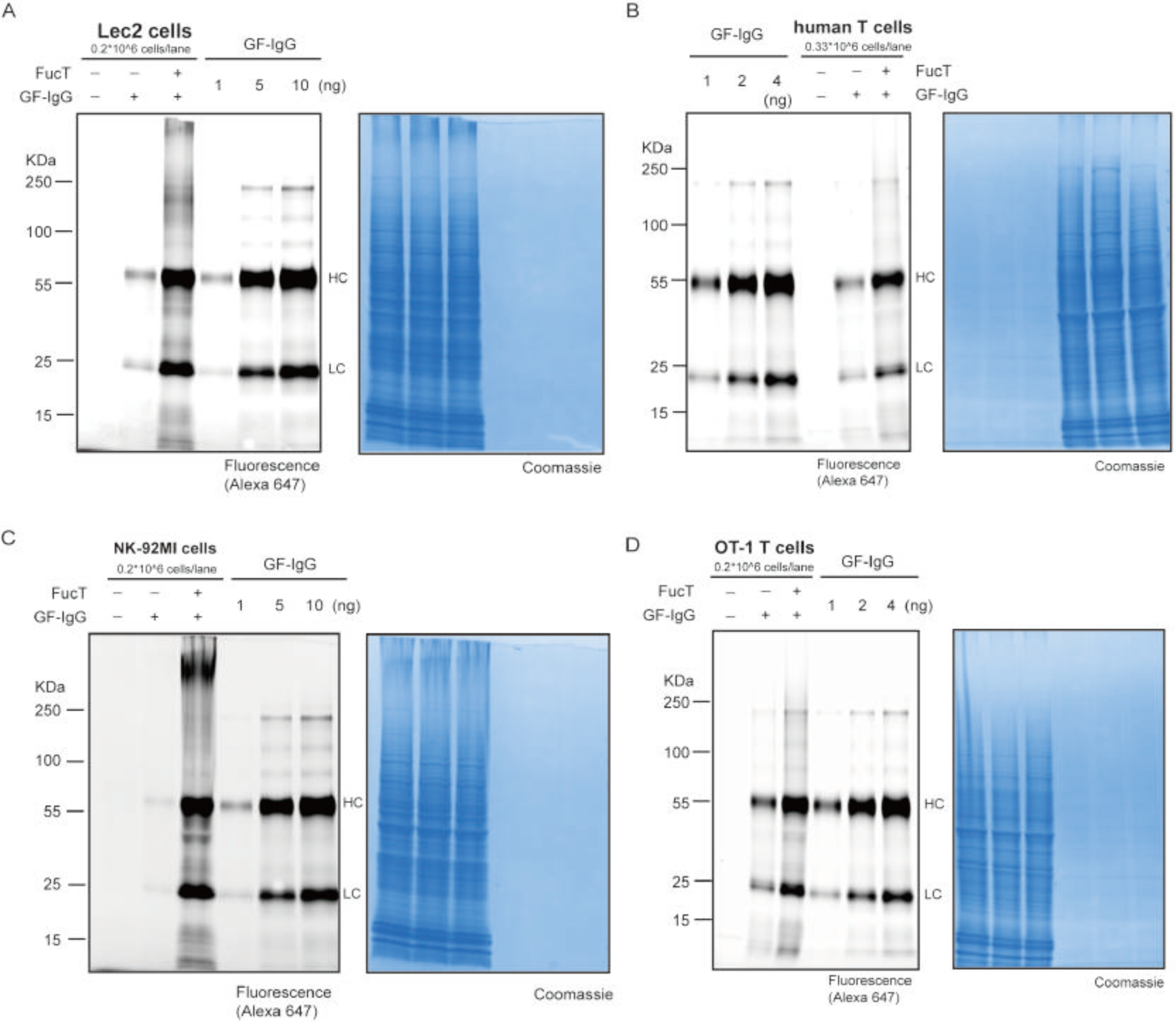
Fluorescent SDS-PAGE gel analysis of IgG labeled cells. GF-IgG molecules modified with Alexa Fluor 647 were used in the experiments for quantifying the number of IgG molecules conjugated to one cell surface. Different amounts of pure GF-IgG-A647 proteins were used as standards in the quantification. At the saturated condition, approximately 2.5×10^5^ IgG molecules were introduced to one Lec2 cell surface (**A**); approximately 2.9×10^4^ IgG molecules were introduced to one primary human T cell surface (**B**); approximately 3×10^5^ IgG molecules were introduced to one NK-92MI cell surface (**C**); approximately 7×10^4^ IgG molecules were introduced to one mouse OT-1 T cell surface (**D**).

**Figure S7.**
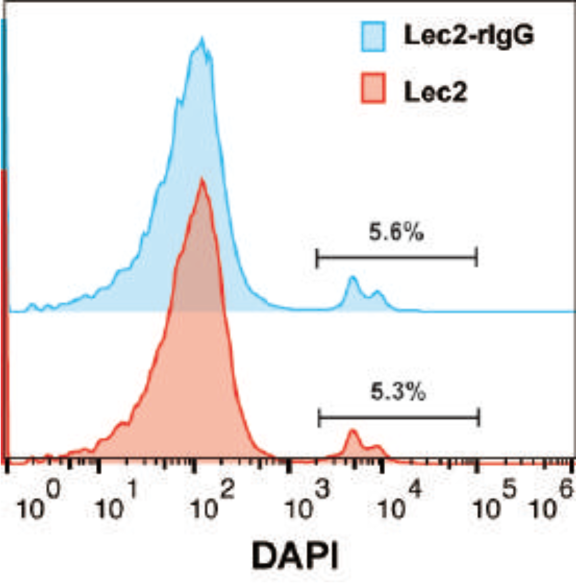
Flow cytometry analysis of the viability of Lec2 cells before and after IgG conjugation.

**Figure S8.**
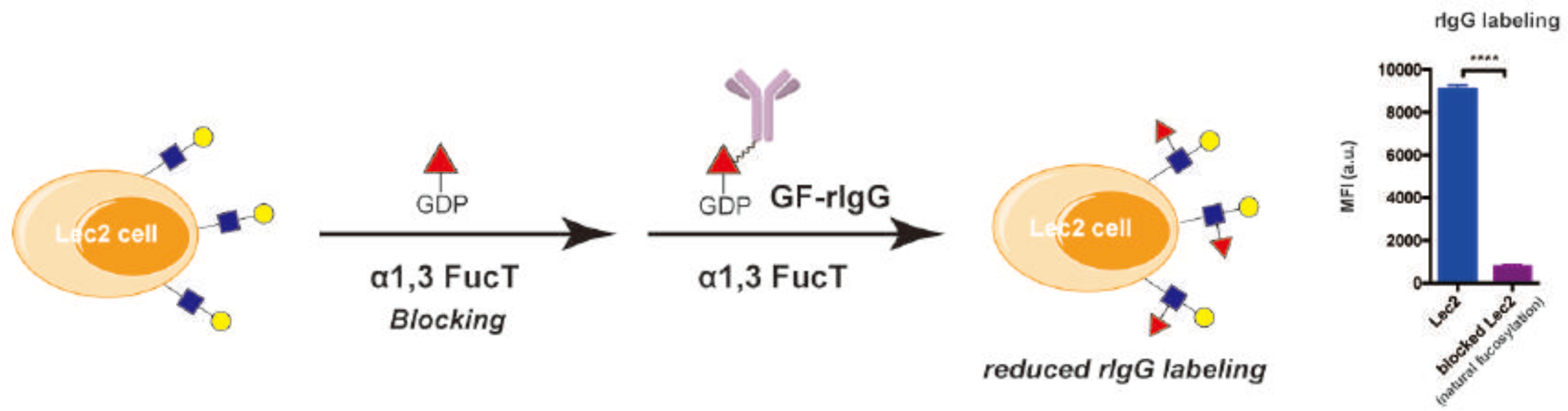
The blockade of LacNAc epitope inhibits FT mediated GF-rIgG transfer to Lec2 cell surface. Lec2 cells were treated with FT and GDP-Fuc first, which could occupy almost all of the fucosylation sites of cell-surface LacNAc. After that, blocked cells were treated with FT and GF-rIgG. rIgG labeling were analyzed by flow cytometry. Compared to direct GF-rIgG transfer, the blocked Lec2 have a significantly reduced rIgG signal. Error bars, mean values ± SD. ^****^P < 0.0001.

**Figure S9.**
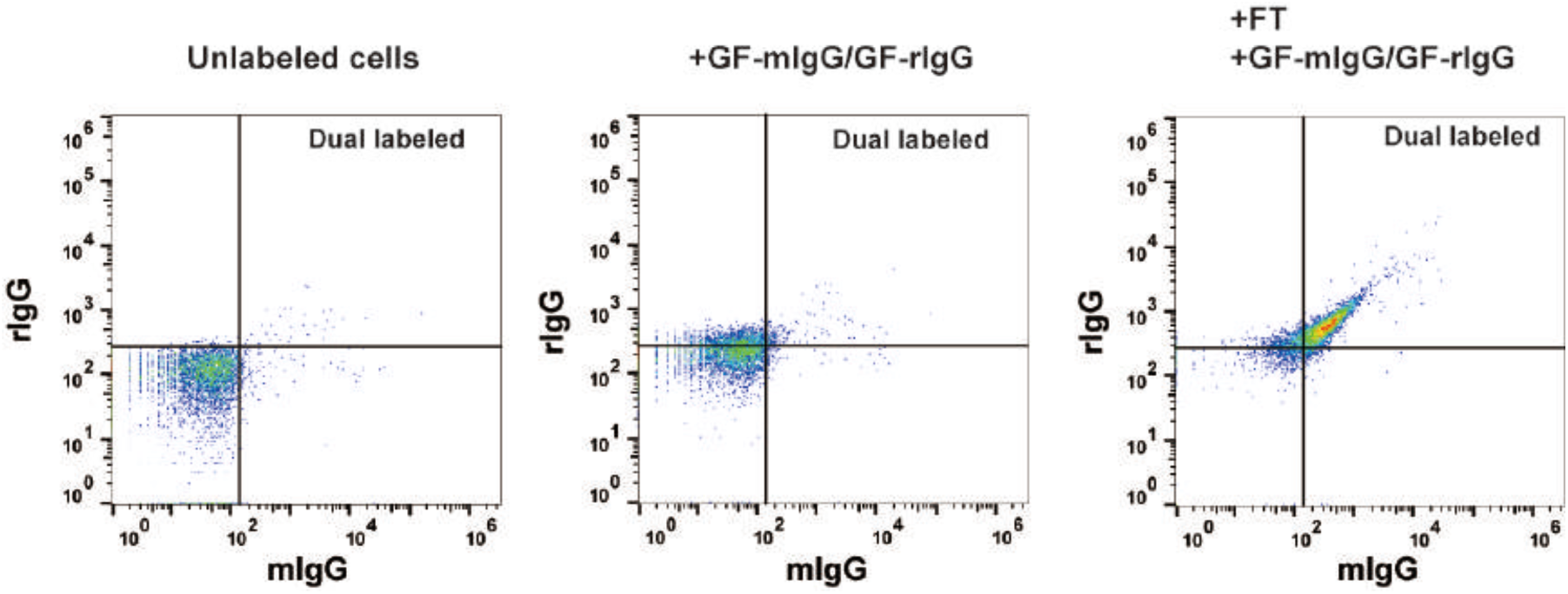
Dual labeling of two different IgG molecules on Lec2 cells. Lec2 cells were treated with GF-mIgG, GF-rIgG and FT. GF-mIgG and GF-rIgG treated was shown as a negative control. The labeled cells were stained with anti-mIgG and anti-rIgG fluorescent antibodies and analyzed by flow cytometry.

**Figure S10.**
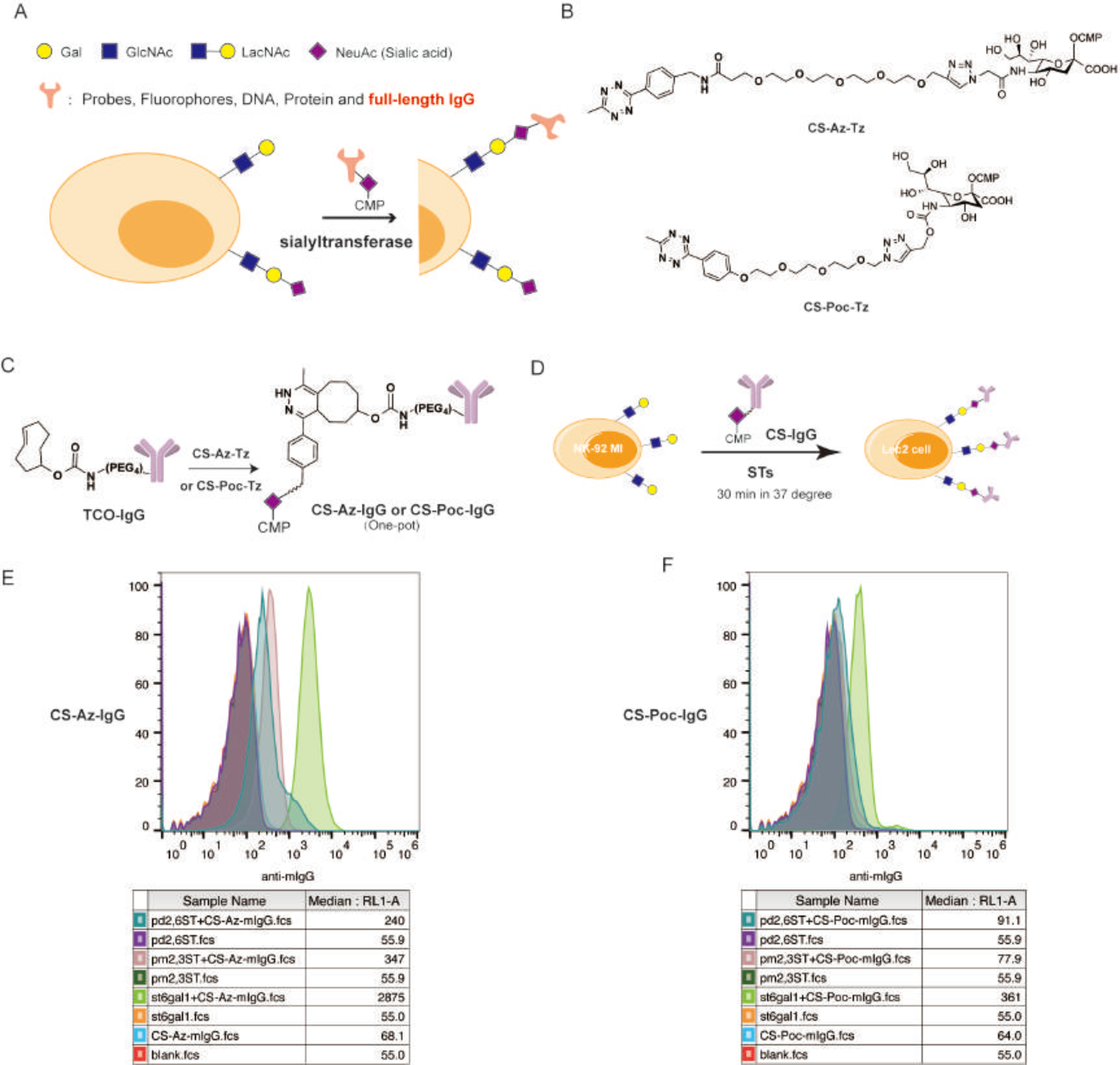
Sialyltransferase enabled labeling of IgG molecules to NK-92MI cell surface. (**A**) Scheme of sialyltransferase enabled enzymatic transferring of functional molecules to cells in one step. (**B**) Chemical structures of CMP-sialic acid-azide-tetrazine (CS-Az-Tz) and CMP-sialic acid-propargyl carbamate-tetrazine (CS-Poc-Tz). (**C**) Schematic representation of the synthesis of a CMP-Sialic acid conjugated IgG (CS-IgG). (**D**) Workflow of the ST-catalyzed transfer of CS-IgG to the surface of NK-92MI cells. (Three different sialyltransferases were tested here). (**E**) Flow cytometry analysis of NK-92MI cells treated with enzyme STs, substrates CS-Az-mIgG, or both. (**F**) Flow cytometry analysis of NK-92MI cells treated with enzyme STs, substrates CS-Poc-mIgG, or both. **Description of this figure**: We first synthesized CMP-sialic acid-azide-tetrazine (CS-Az-Tz) and CMP-sialic acid-propargyl carbamate-tetrazine (CS-Poc-Tz) using one-pot click chemistry (B). Then, mouse IgG2a antibodies bearing TCO moieties were reacted with CS-Az-Tz or CS-Poc-Tz via the inverse electron-demand Diels–Alder reaction (IEDDA) to generate CMP-Sialic acid-conjugated IgG molecules (CS-IgG) (C). The one-pot product of CMP-Sialic acid conjugated mouse IgG (CS-mIgG) was then incubated with NK-92 MI cells that express abundant terminal LacNAc units in the presence of sialyltransferases (STs) (D). We tested three STs, including recombinant ST6Gal1, *Pasteurella multocida* α(2,3) sialyltransferase M144D mutant (Pm2,3ST-M144D), *Photobacterium damsel* α(2,6) sialyltransferase (Pd2,6ST). According to the results, all of these three enzymes could transfer CS-mIgG to cell surface (E and F). In general, CS-Az-IgG is a more favorable substrate of these STs than CS-Poc-IgG. ST6Gal1 is the most efficient ST in these three enzymes (E and F). Taken together, ST6Gal1 and CS-Az-IgG is the best pair to transfer IgG to cell surface through sialylation (Figure E). This example demonstrates cell surface engineering through glycan to install biomacromolecules could also be achieved using sialyltranferases.

**Figure S11.**
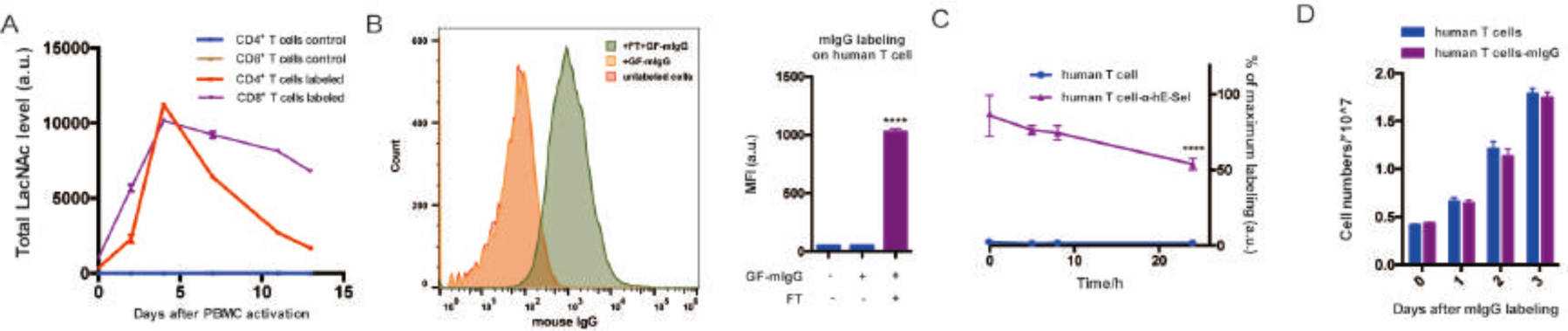
Transferring IgG molecules onto human T cells. (**A**) LacNAc on Human PBMC was labeled by GF-Biotin on different days after activation. Labeled cells were stained with streptavidin-APC, anti-CD4 and anti-CD8 fluorescent antibodies, and analyzed by flow cytometry. (**B**) Human T cells were treated with FT and GF-mIgG, or GF-mIgG alone, or untreated. After 15 min of labeling, cells were stained with anti-mIgG fluorescent antibody and analyzed by flow cytometry. (**C**) Human T cells labeled with mIgG were stained with anti-mIgG fluorescent antibody and analyzed by flow cytometry at different time points after labeling. (**D**) Human T cells with or without mIgG labeling were cultured in T cell media and live cells in each group were counted on different days after labeling. Error bars, mean values ± SD. In all figures: ****P < 0.0001.

**Figure S12.**
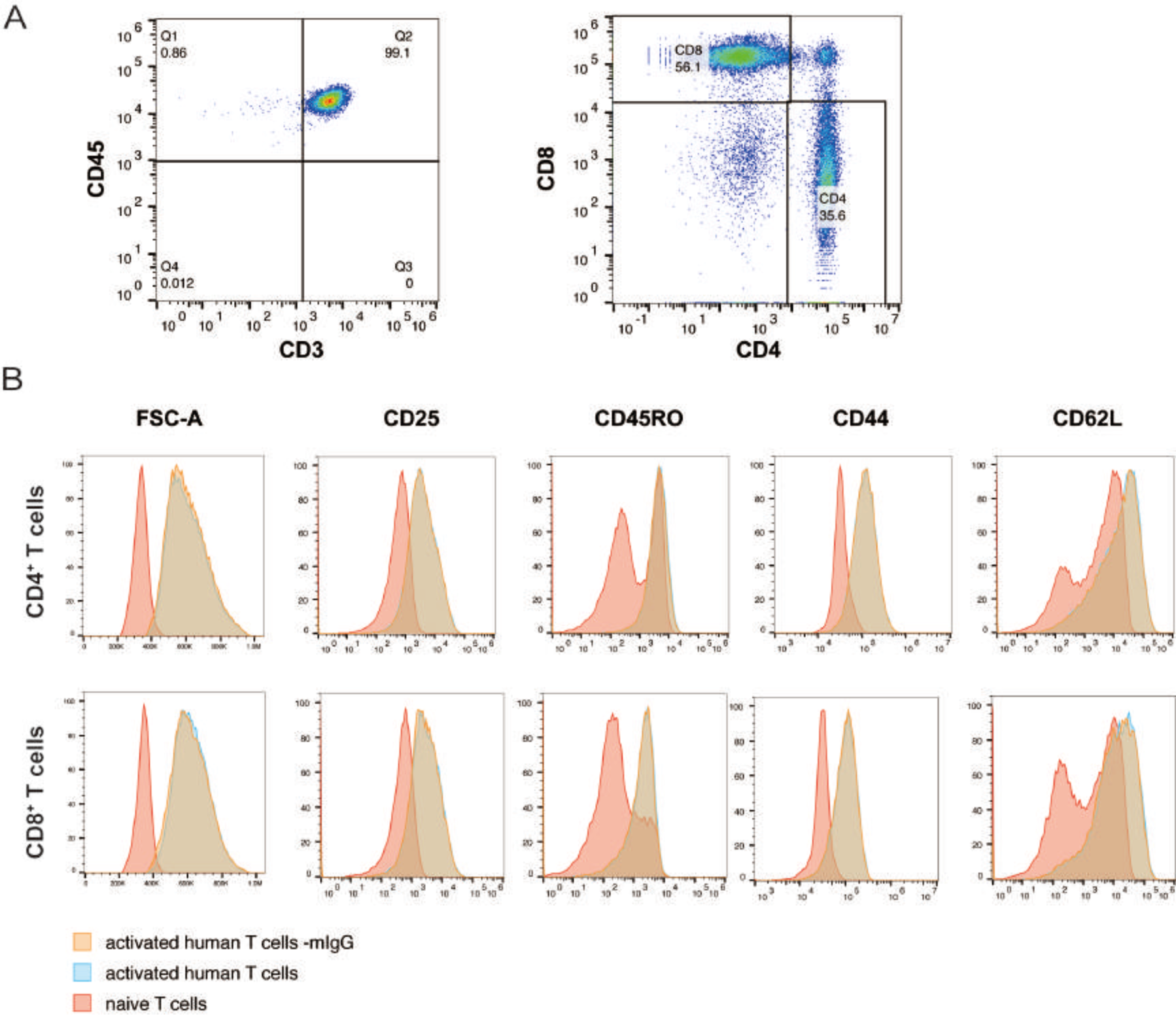
Phenotype of human T cells before and after mIgG labeling. (**A**) Human PBMC were activated by anti-CD3/CD28 antibody and expanded *in vitro* for about two weeks. The cells were stained with anti-CD3, anti-CD45, anti-CD4 and anti-CD8 fluorenscent antibodies, and analyzed by flow cytometry. (**B**) Activated human T cells were treated with GF-IgG and FT. Labeled or unlabeled cells were stained with anti-CD4, anti-CD8, anti-CD25, anti-CD45RO, anti-CD44 and anti-CD62L fluorescent antibodies and analyzed by flow cytometry. Naïve T cells were used as control.

**Figure S13.**
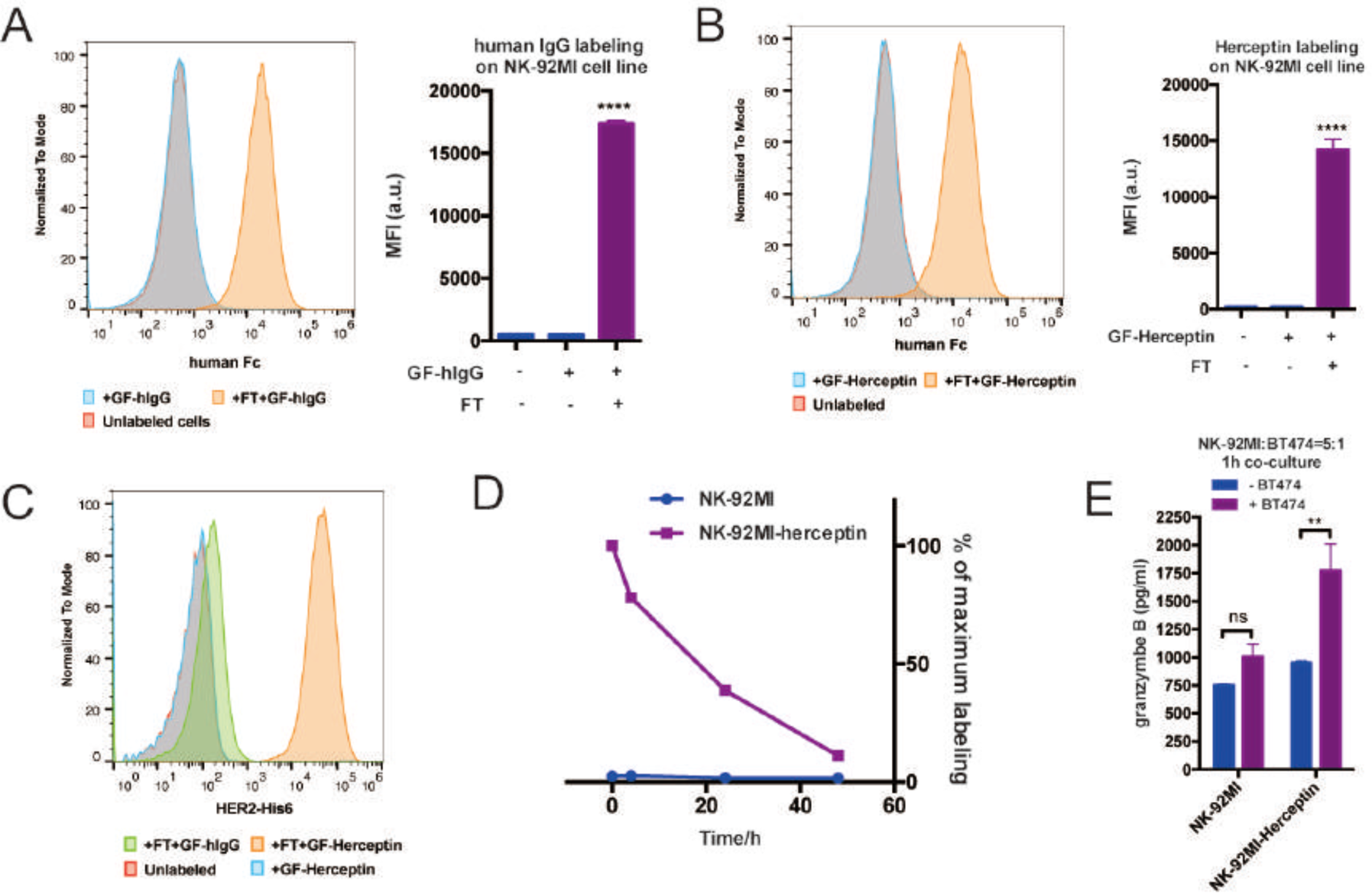
Transferring IgG molecules onto NK-92MI cells. (**A**) NK-92MI cells were treated with GF-hIgG and FT, or GF-hIgG alone, or untreated. The cells were stained with anti-hFc fluorescent antibody and analyzed by flow cytometry. (**B**) NK-92MI cells were treated with GF-Herceptin and FT, or GF-Herceptin alone, or untreated. The cells were stained with anti-hFc fluorescent antibody and analyzed by flow cytometry. (**C**) NK-92MI cells with different modifications were then incubated with HER2-His6 protein and then stained with anti-His6 fluorescent antibody for FACS analysis. (**D**) NK-92MI cells conjugated with Herceptin were stained with anti-hFc fluorescent antibody and analyzed by flow cytometry at different time points post labeling. (**E**) NK-92MI cells conjugated with or without Herceptin were co-cultured with BT474 at a ratio of 5:1 (Effector: Target) for 1h. Granzyme B secretion in culture supernatant was analyzed by ELISA. Error bars, mean values ± SD. In all figures: ns, P >0.05; **P < 0.01.

**Figure S14.**
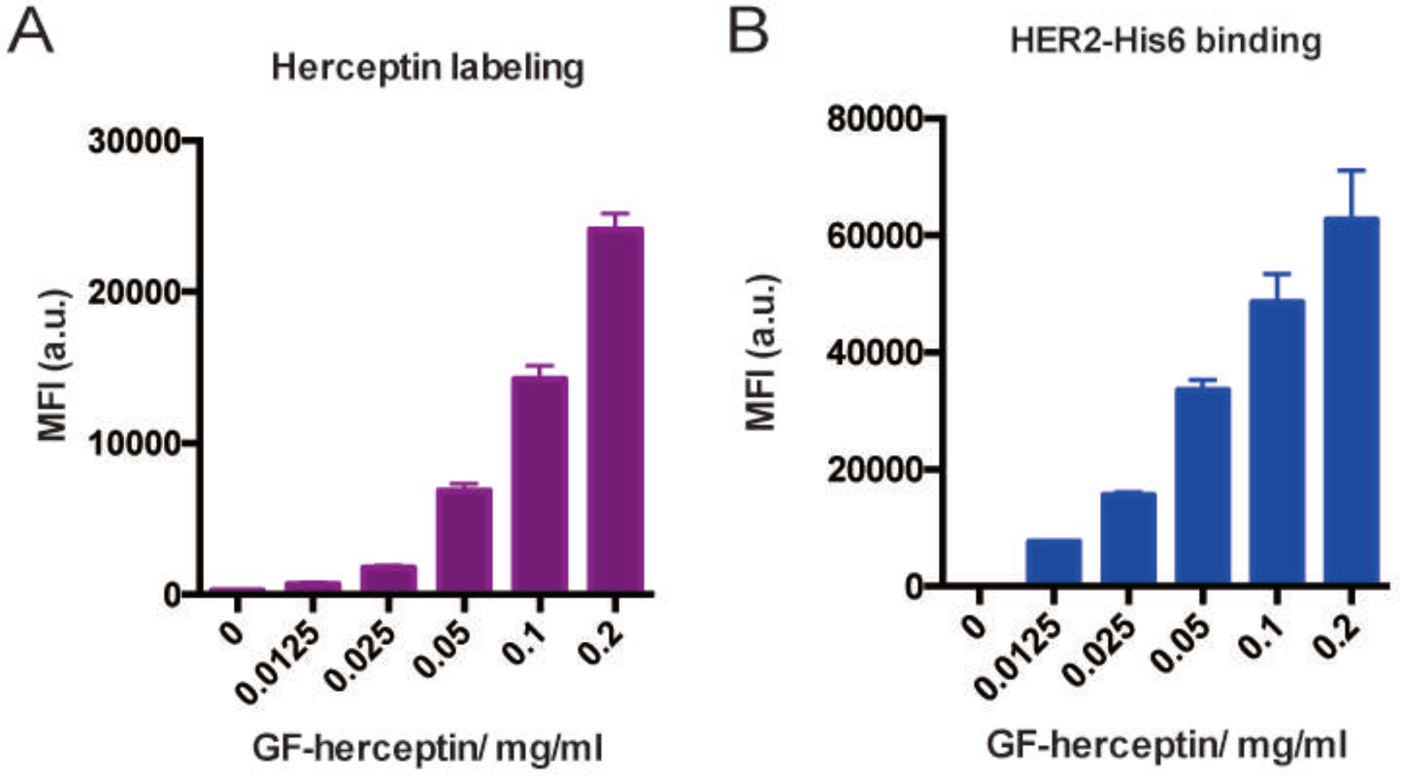
Titration of GF-Herceptin concentrations in the enzymatic transfer. NK-92MI cells were treated with different concentrations of GF-Herceptin in a standard labeling condition. Labeled cells were directly stained with anti-hFc fluorescent antibody (**A**), or incubated with HER2-His6 and then stained with anti-His6 fluorescent antibody (**B**) before flow cytometry analysis.

**Figure S15.**
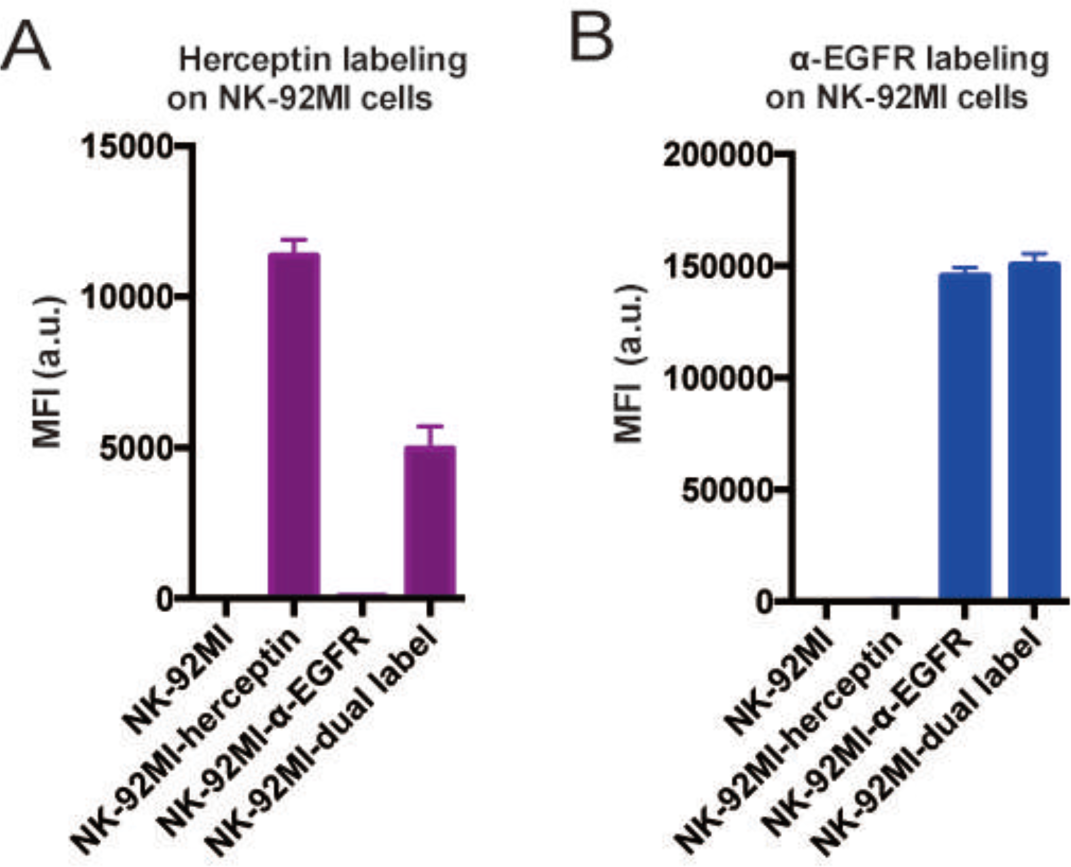
Dual antibody labeling on NK-92MI cells. NK-92MI cells were treated with FT and GF-Herceptin, and then treated with FT and GF-α-EGFR. Labeled cells were stained with anti-hFc fluorescent antibody (**A**), or stained with anti-mIgG fluorescent antibody (**B**) before flow cytometry analysis.

**Figure S16.**
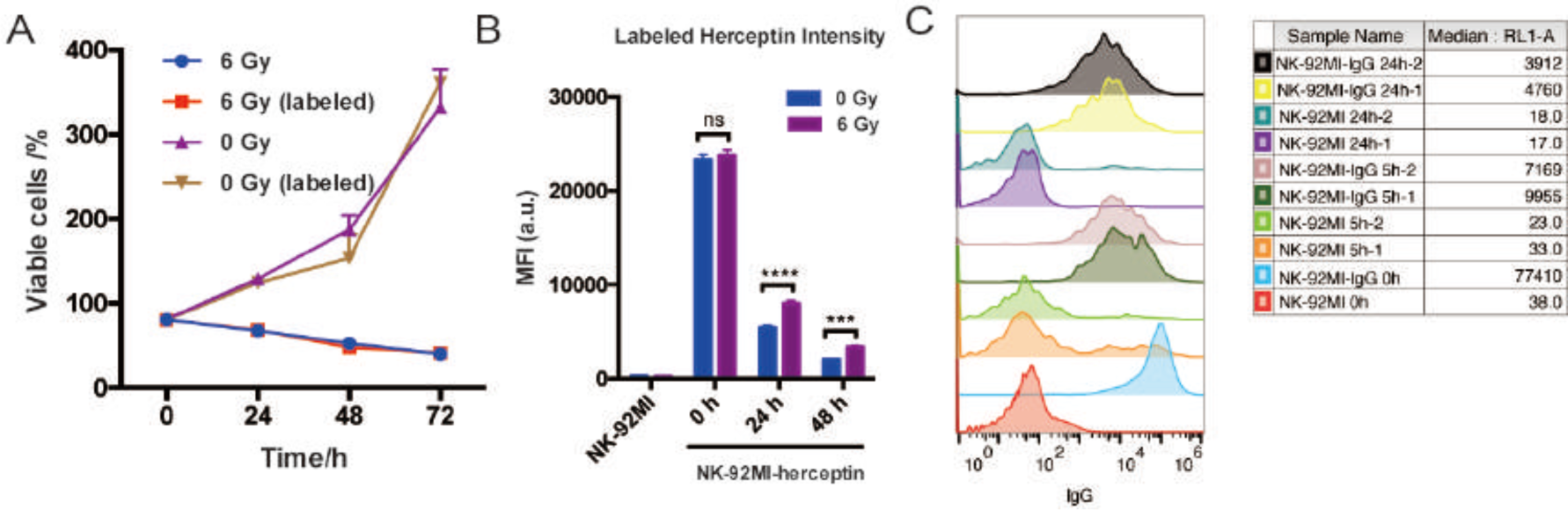
Proliferation and stability of Herceptin-NK-92MI conjugates with or without γ-irradiation. NK-92MI cells were received with or without 6 Gy gamma irradiations before enzymatic reaction. Labeled and unlabeled NK-92MI cells were cultured for another three days. (**A**) Viable cells were counted by flow cytometry at different time points after irradiation and labeling. (**B**) Herceptin labeled cells were also stained with anti-hFc fluorescent antibody and analyzed by flow cytometry at different time points post labeling. Irradiated cells have a slower decay of Herceptin labeling. (**C**) Herceptin conjugated to NK-92MI cell surface was tracked after these cells (non-irradiated) were injected into NSG mice (*i.v.*). NK-92MI cells were labeled using cell tracker (orange) before the Herceptin conjugation. Blood were collected after 5 hours or 24hours after injection. The red blood cells were lysed before all the other cells were analyzed directly through FACS. Error bars, mean values ± SD. In all figures: ns, P >0.05; ***P < 0.001; ****P < 0.0001.

**Figure S17.**
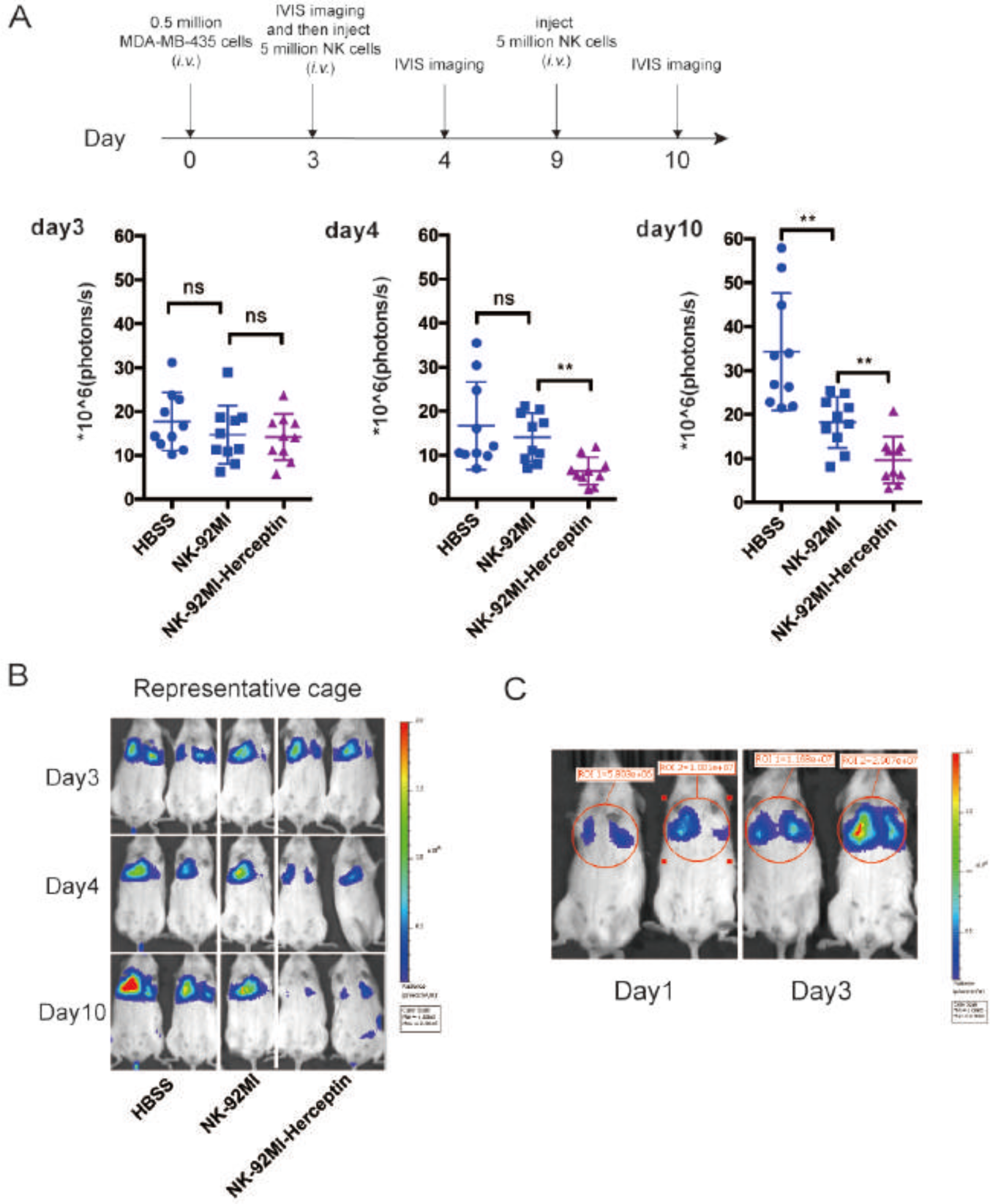
Herceptin-NK-92MI conjugate shows enhanced antitumor activity in established lung metastasis mice model. (**A**) On day 0, NSG mice were injected intravenously with 0.5 million MDA-MB-435/HER2+/F-luc cells. Three days later, the animals were imaged in IVIS system to detect bioluminescent signals for the confirmation of tumor formation and then treated by *i.v.* injection of 5 million NK-92MI or Herceptin-NK-92MI cells. The control mice received HBSS. The animals were imaged again to see the efficacy of NK cell treatment on day 4 and day 10. The animals were received one more treatment of 5 million NK-92MI or Herceptin-NK-92MI cells (*i.v.* injection) on day 9. The sizes of the tumors of the mice and mean values ± SD are shown; n = 10. (**B**) Representative images also are shown. (**C**) Two mice were imaged on day 1 and day3 to show a 3fold increase in bioluminescence, which confirms that tumors are growing after inoculation. In all figures, ns, P > 0.05; ^**^P < 0.01.

**Figure S18.**
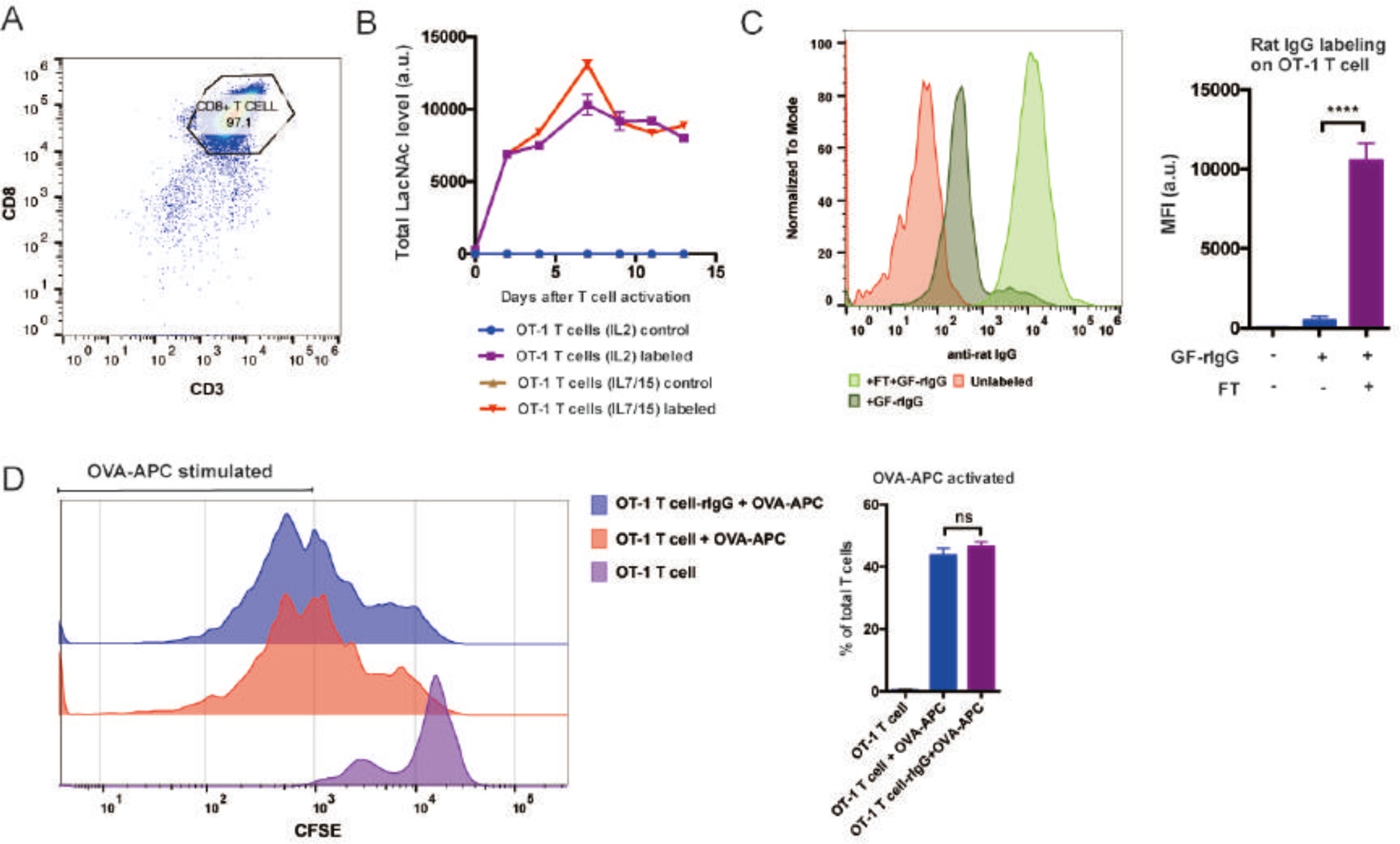
Enzymatic transfer of rIgG to OT-1 CD8+ T cells. (**A-B**) Spleenocytes from OT-1 mice were activated by OVA peptides and *in vitro* expanded by adding IL2 or IL7/IL15. After four days expansion, the cells were stained with anti-CD3 and anti-CD8 fluorescent antibodies and analyzed by flow cytometry (A). The cells were also treated with GF-Al-Biotin and FT, and then stained with streptavidin-APC and analyzed by flow cytometry to track LacNAc level on different days after T cell activation (B). (**C**) OT-1 T cells were treated with GF-rIgG and FT, or GF-rIgG alone, or untreated. Then the cells were stained with anti-rIgG fluorescent antibody and analyzed by flow cytometry. (**D**) OT-1+/-CD45.1+/-T cells were first labeled with CFSE and then treated with GF-rIgG and FucT, or untreated. After fucosylation, the cells were co-cultured with OVA peptide pulsed WT B6 spleenocytes for 48h and T cell proliferation was analyzed by flow cytometry. Error bars, mean values ± SD. In all figures: ns, P >0.05; ****P < 0.0001.

**Figure S19.**
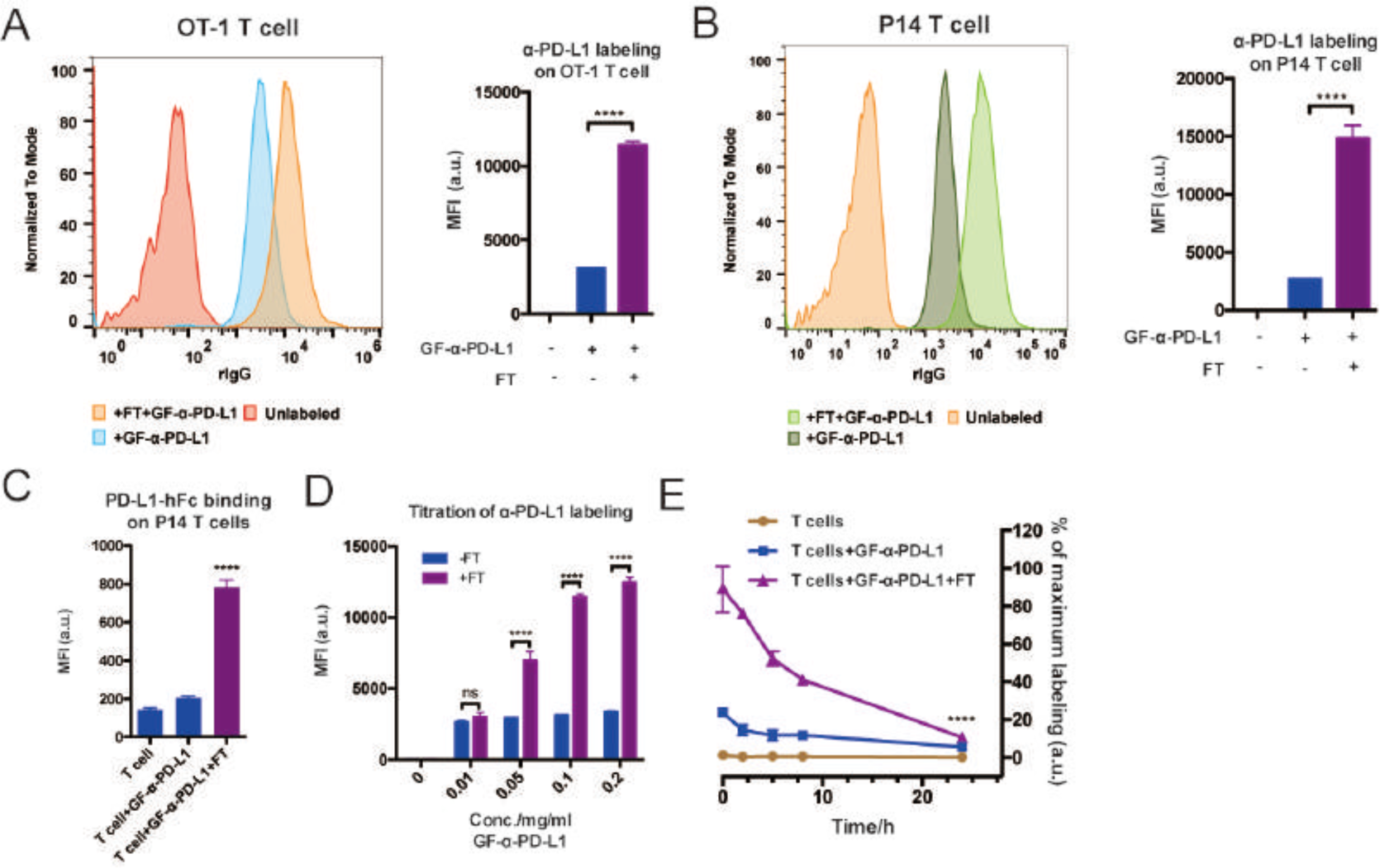
Transferring α-PD-L1 onto OT-1 CD8+ T cells. (**A**) OT-1 CD8+ T cells were treated with GF-α-PD-L1 and FT, or GF-α-PD-L1 alone, or untreated. The cells were then stained with anti-rIgG fluorescent antibody and analyzed by flow cytometry. (**B-C**) CD8+ T cells from P14 mice were treated with GF-α-PD-L1 and FT, or GF-α-PD-L1 alone, or untreated. The cells were stained with anti-rIgG fluorescent antibody (B), or incubated with PD-L1-hFc first and then stained with anti-hFc fluorescent antibody (C). After staining, these cells were analyzed by flow cytometry. (**D**) OT-1 T cells were labeled under different concentrations of GF-α-PD-L1 with or without FT. After labeling, cells were stained with anti-rIgG fluorescent antibody and analyzed by flow cytometry. (**E**) OT-1 T cells were treated with GF-α-PD-L1 and FT, or GF-α-PD-L1 alone, or untreated. The cells were then stained with anti-rIgG fluorescent antibody and analyzed by flow cytometry at different time points after reaction. Error bars, mean values ± SD. In all figures: ns, P >0.05; ****P < 0.0001.

**Figure S20.**
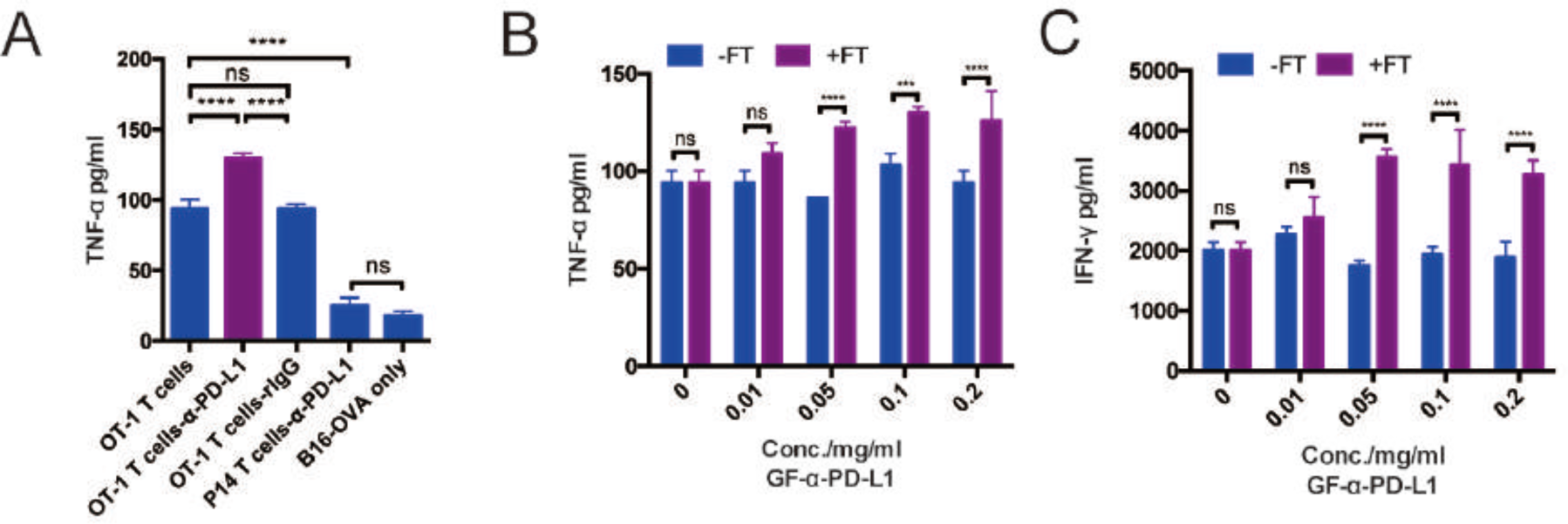
Transferring α-PD-L1 onto OT-1 T cell surface enhance the cytokine secretion during killing cancer cells. (**A**) OT-1 T cells and P14 T cells were treated with GF-α-PD-L1 and FT, or GF-rIgG and FT, or untreated. T cells were then co-cultured with B16-OVA cells for 9 hours. TNF-α concentrations in culture supernatant were quantified via ELISA kit. (**B-C**) OT-1 T cells were treated under different concentrations of GF-α-PD-L1 with or without FT, and then co-cultured with B16-OVA cells for 9 hours. TNF-α and IFN-γ in culture supernatant were analyzed by ELISA. Error bars, mean values ± SD. In all figures: ns, P >0.05; ***P < 0.001; ****P < 0.0001.

**Figure S21.**
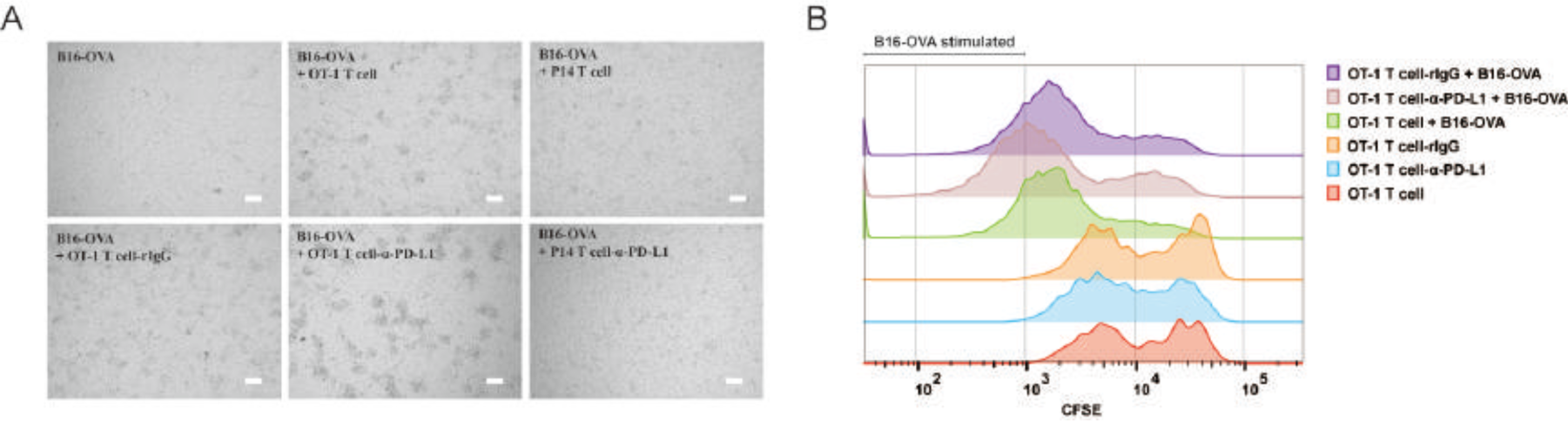
Transferring α-PD-L1 onto T cell surface increases the size of T-cell clustering and T cell proliferation during killing. (**A**) OT-1 T cells and P14 T cells were treated with GF-α-PD-L1 and FT, or GF-rIgG and FT, or untreated. The cells were then co-cultured with B16-OVA cells for 20 hours. T cell cluster during killing were imaged using microscope. Scale bar: 50 μm. (**B**) OT-1 T cells were stained with CFSE and treated with GF-α-PD-L1 and FT, or GF-rIgG and FT, or untreated. Then the cells were cultured with or without B16-OVA cells for 72 hours. Proliferations of OT-1 T cells were analyzed through CFSE signal dilution.

## Spectra

### 1. Spectra of GF-Al-Tz

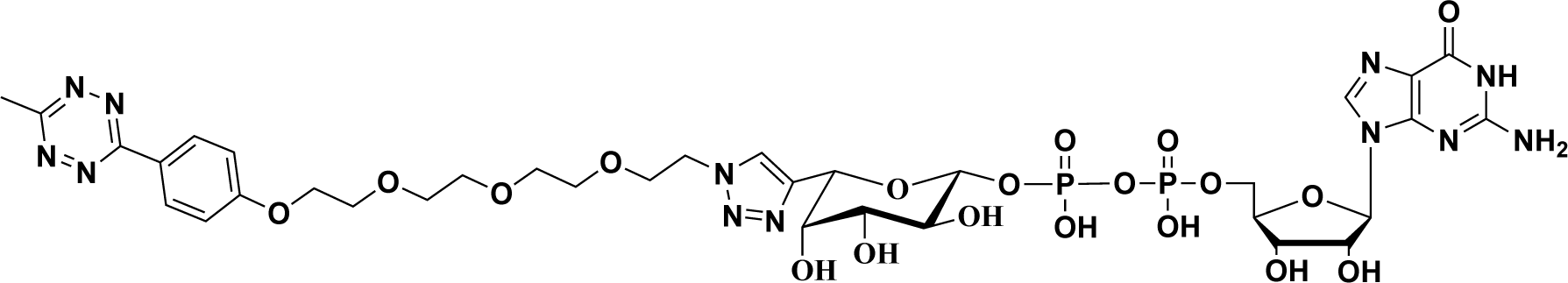

TLC

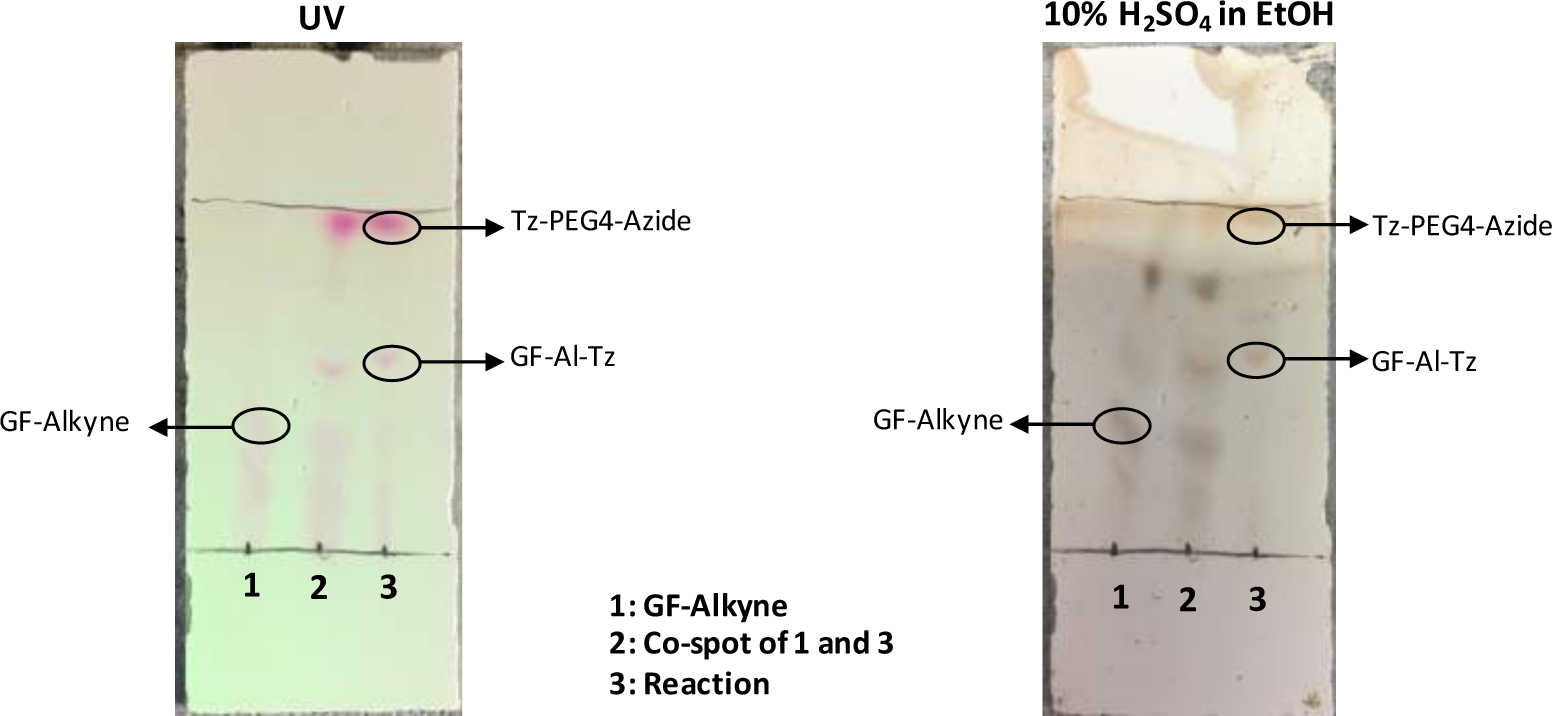

ESI-TOF MS

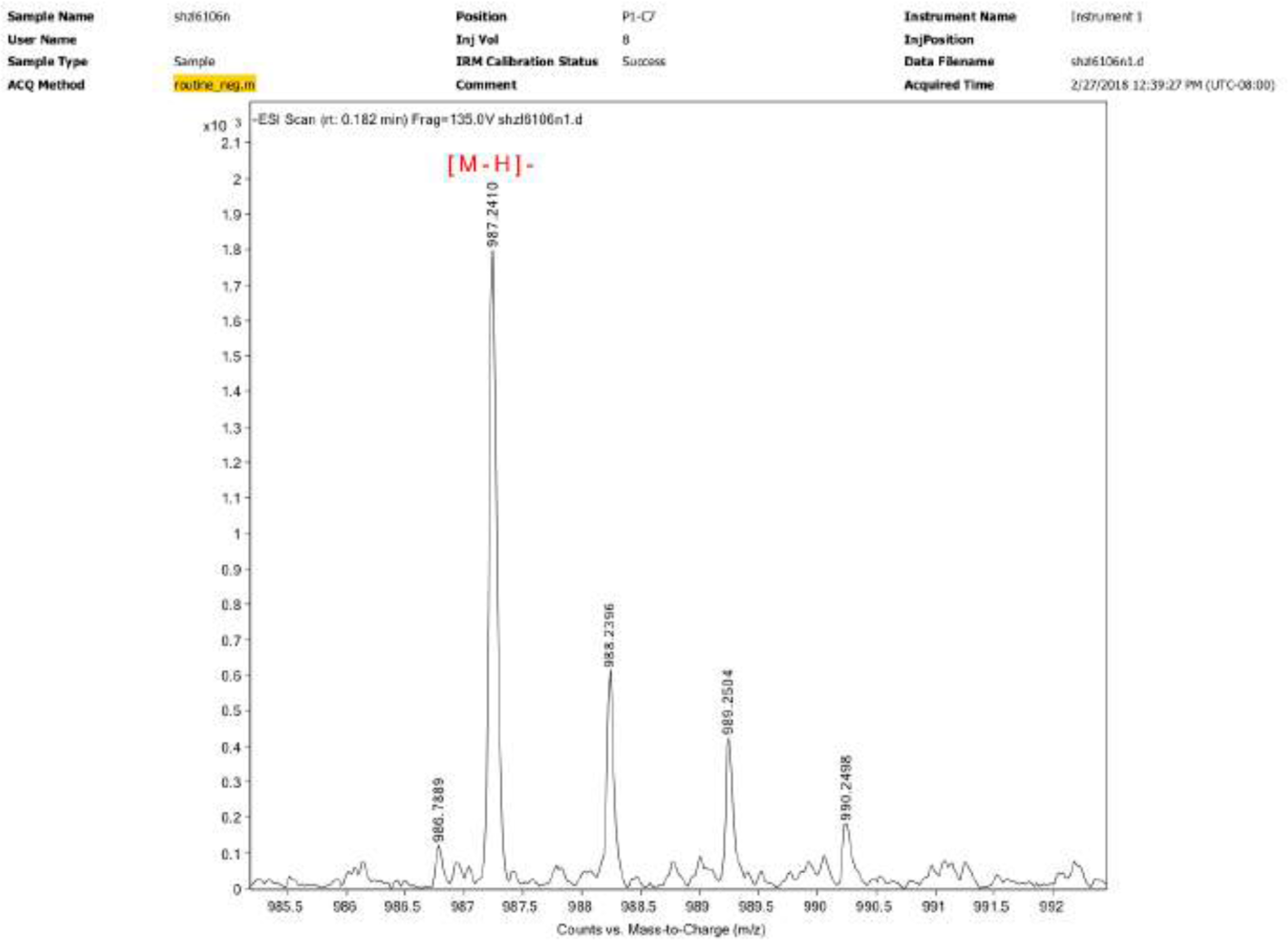

^1^H NMR

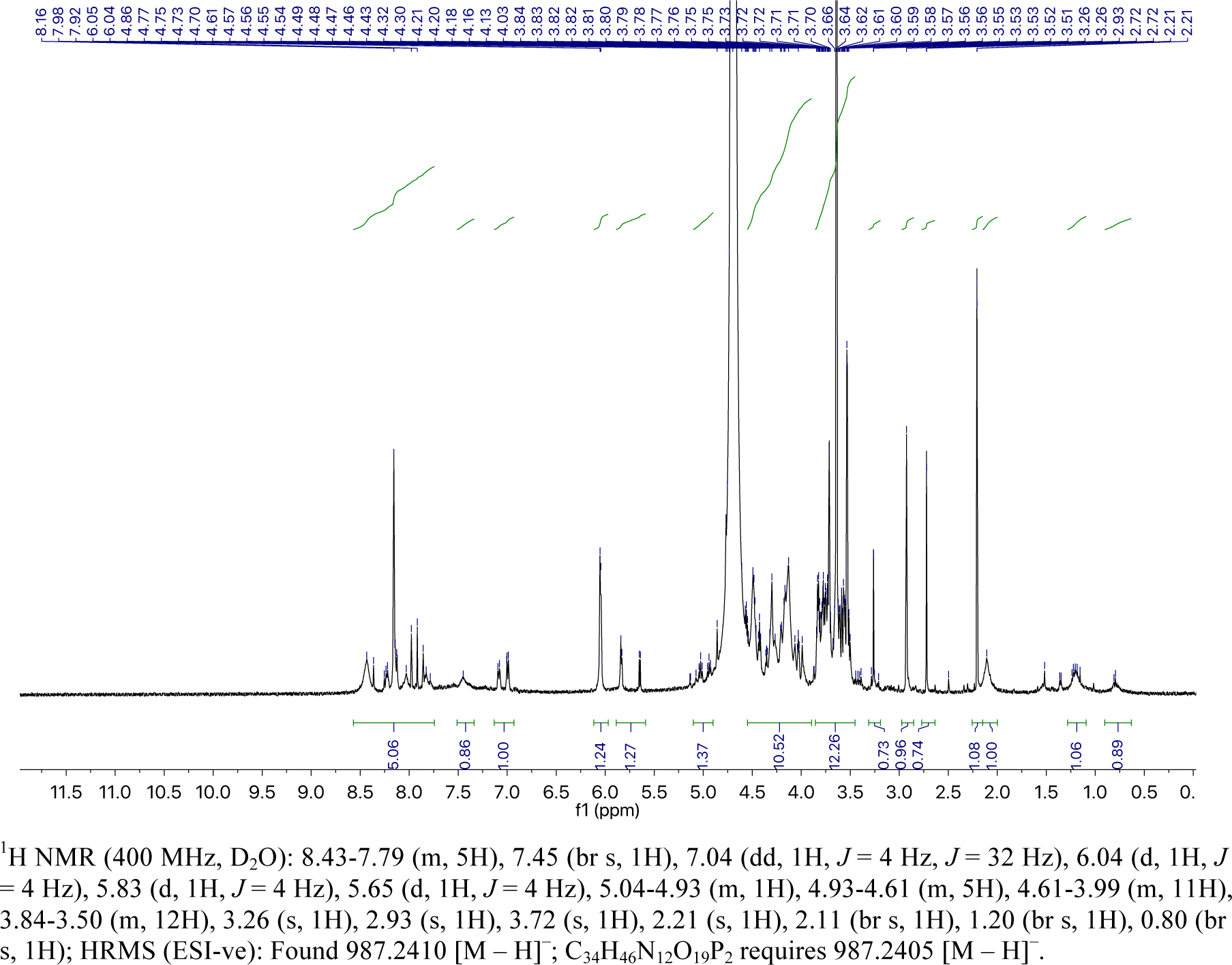

### 2. Spectra of GF-Az-Tz

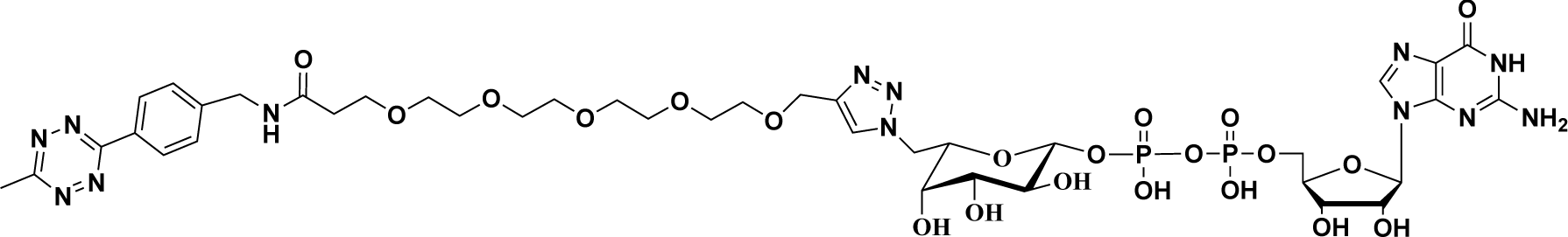

TLC

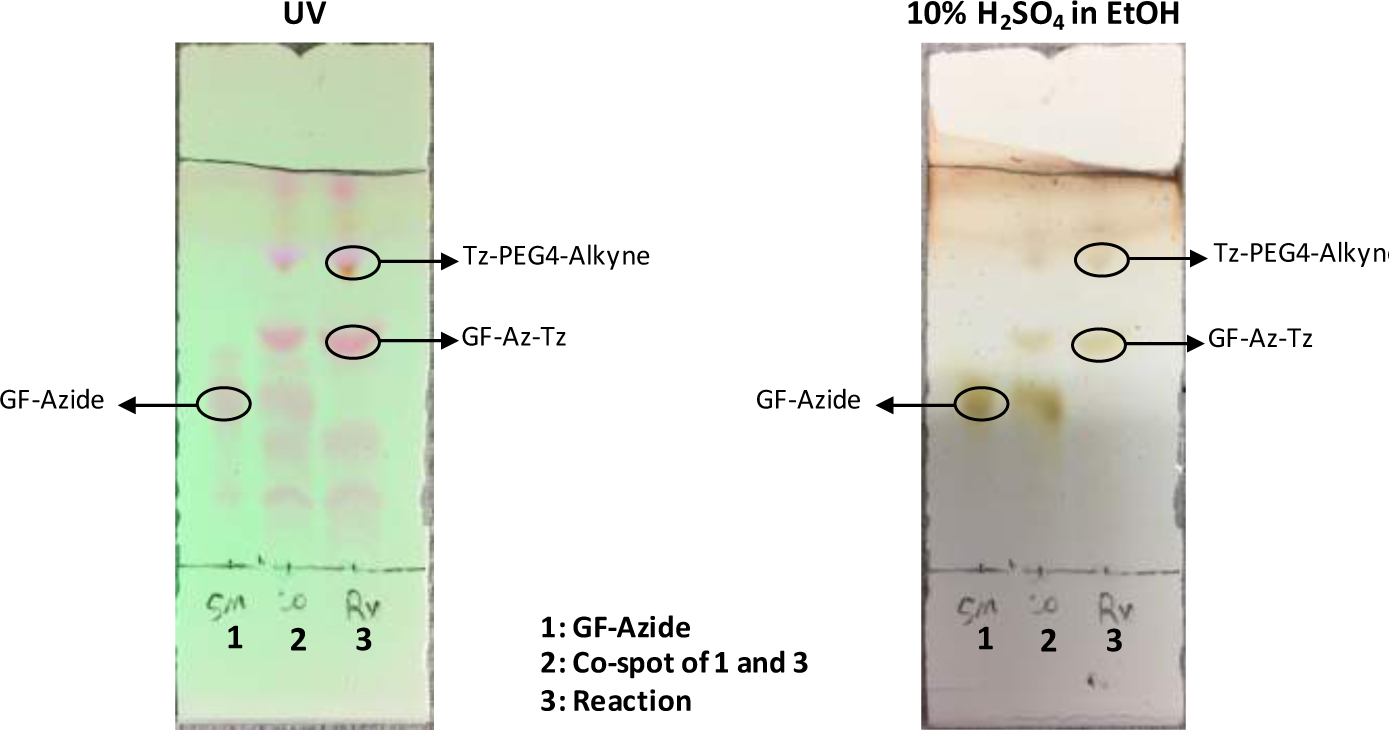

ESI-TOF MS

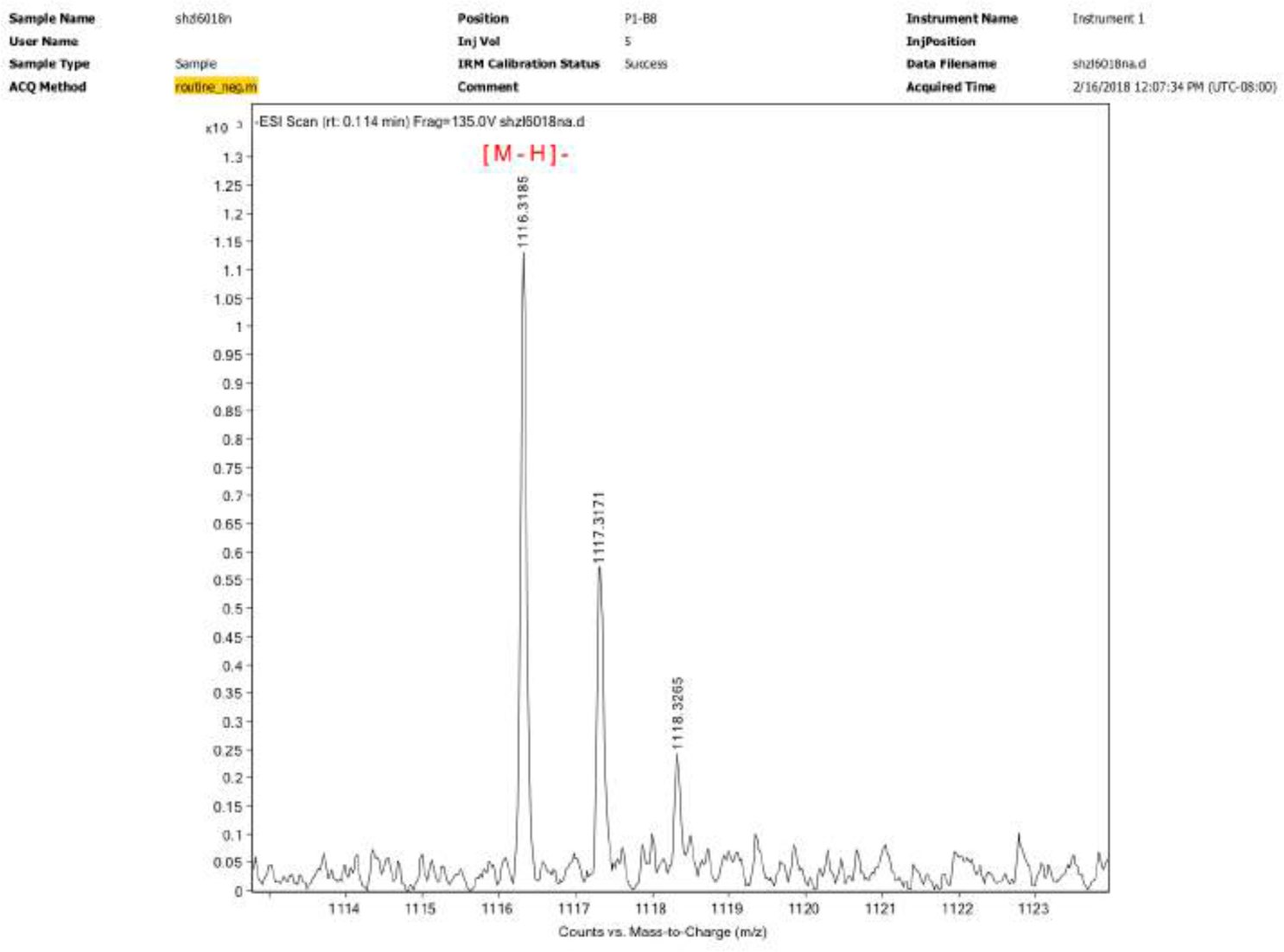

^1^H NMR

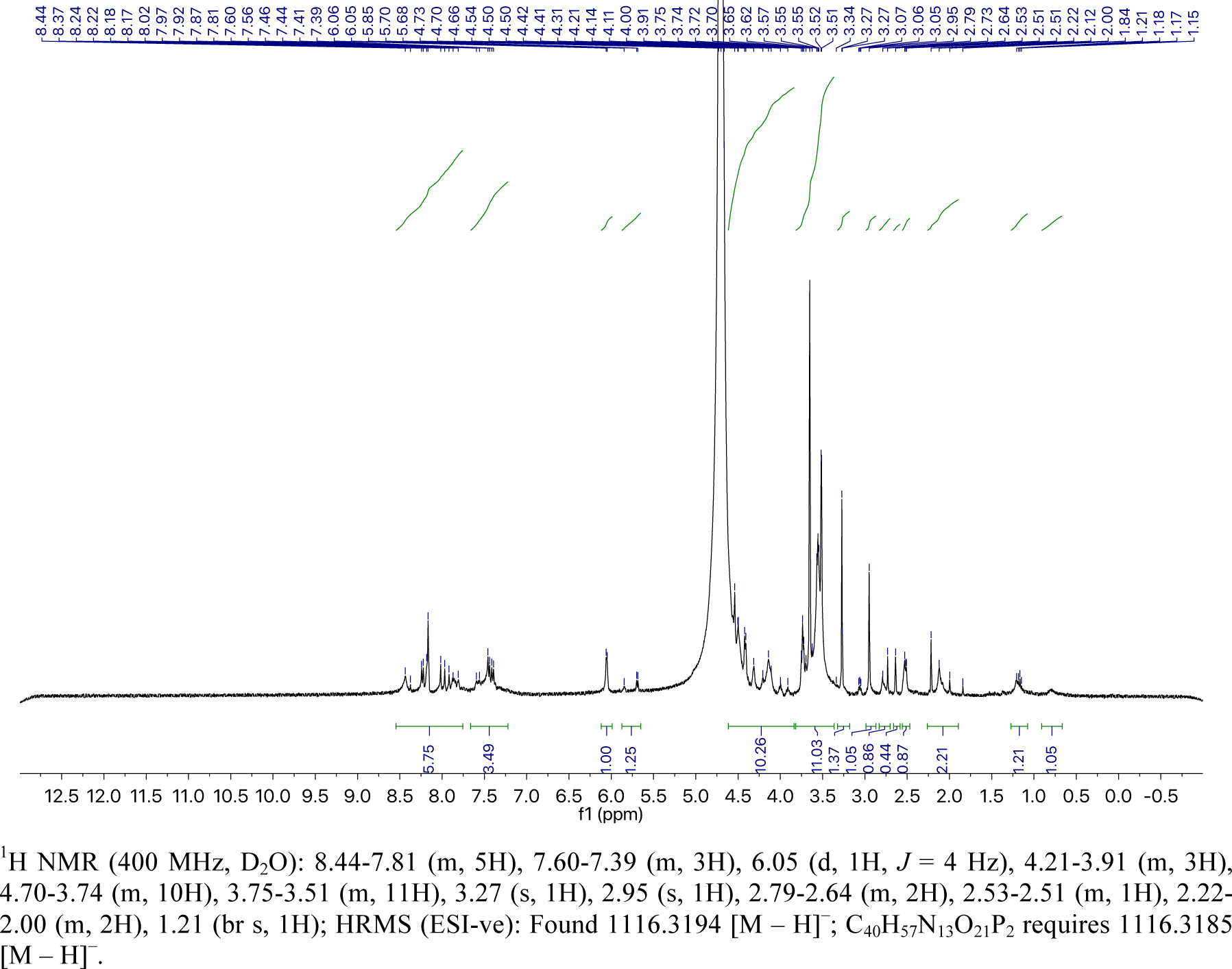

## Reference

(1) Rosenberg, S. A.; Restifo, N. P. Science 2015, 348, 62.

(2) Prasad, V. Nat. Rev. Clin. Oncol. 2017.

(3) Cornetta, K.; Pollok, K. E.;Miller, A. D. Cold Spring Harbor Protocols 2008, 2008, pdb.prot4884.

(4) Dupré, L.; Trifari, S.; Follenzi, A.; Marangoni, F.; Lain de, Lera T.; Bernad, A.; Martino, S.; Tsuchiya, S.; Bordignon, C.; Naldini, L.; Aiuti, A.; Roncarolo, M.-G. Mol. Ther. 2004, 10, 903.

(5) Shearer, R. F.; Saunders, D. N. Genes Cells 2015, 20, 1.

(6) Stephan, M. T.; Irvine, D. J. Nano Today 2011, 6, 309.

(7) Swartz, M. A.; Hirosue, S.; Hubbell, J. A. Sci. Transl. Med. 2012, 4, 148rv9.

(8) Griffin, Matthew E.; Hsieh-Wilson, Linda C. Cell Chem. Biol. 2016, 23, 108.

(9) Hudak, Jason E.; Bertozzi, Carolyn R. Chem. Biol. 2014, 21, 16.

(10) Bi, X.; Yin, J.; Chen Guanbang, A.; Liu, C.-F. Chemistry – A European Journal 2018, 10.1002/chem.201705049.

(11) Pishesha, N.; Bilate, A. M.; Wibowo, M. C.; Huang, N.-J.; Li, Z.; Deshycka, R.; Bousbaine, D.; Li, H.; Patterson, H. C.; Dougan, S. K.; Maruyama, T.; Lodish, H. F.; Ploegh, H. L. Proc. Natl. Acad. Sci. USA 2017, 114, 3157.

(12) Parmar, S.; Liu, X.; Najjar, A.; Shah, N.; Yang, H.; Yvon, E.; Rezvani, K.; McNiece, I.; Zweidler-McKay, P.; Miller, L.; Wolpe, S.; Blazar, B. R.; Shpall, E. J. Blood 2015, 125, 1502.

(13) Sackstein, R.; Merzaban, J. S.; Cain, D. W.; Dagia, N. M.; Spencer, J. A.; Lin, C. P.; Wohlgemuth, R. Nat. Med. 2008, 14, 181.

(14) Srivastava, G.; Kaur, K. J.; Hindsgaul, O.; Palcic, M. M. J. Biol. Chem. 1992, 267, 22356.

(15) Zheng, T.; Jiang, H.; Gros, M.; Soriano del Amo, D.;Sundaram, S.; Lauvau, G.; Marlow, F.; Liu, Y.; Stanley, P.; Wu, P. Angew. Chem. Int. Ed. 2011, 123, 4199.

(16) Capicciotti, C. J.; Zong, C.; Sheikh, M. O.; Sun, T.; Wells, L.; Boons, G.-J. J. Am. Chem. Soc. 2017, 139, 13342.

(17) Wang, W.; Hu, T.; Frantom, P. A.; Zheng, T.; Gerwe, B.; del Amo, D. S.; Garret, S.; Seidel, R. D.; Wu, P. Proc. Natl. Acad. Sci. USA 2009, 106, 16096.

(18) Rostovtsev, V. V.; Green, L. G.; Fokin, V. V.; Sharpless, K. B. Angew. Chem. Int. Ed. 2002, 41, 2596.

(19) Tornøe, C. W.; Christensen, C.; Meldal, M. J. Org. Chem. 2002, 67, 3057.

(20) Besanceney-Webler, C.; Jiang, H.; Zheng, T.; Feng, L.; Soriano del Amo, D.; Wang, W.; Klivansky, L. M.; Marlow, F. L.; Liu, Y.; Wu, P. Angew. Chem. Int. Ed. 2011, 50, 8051.

(21) Swee, L. K.; Lourido, S.; Bell, G. W.; Ingram, J. R.; Ploegh, H. L. ACS Chem. Biol. 2015, 10, 460.

(22) Giorgi, M. E.; Agusti, R.; de Lederkremer, R. M. Beilstein Journal of Organic Chemistry 2014, 10, 1433.

(23) Rahim, M. K.; Kota, R.; Haun, J. B. Bioconjug. Chem. 2015, 26, 352.

(24) Oliveira, B. L.; Guo, Z.; Bernardes, G. J. L. Chem. Soc. Rev. 2017, 46, 4895.

(25) Morvan, M. G.; Lanier, L. L. Nat. Rev. Cancer 2015, 16, 7.

(26) Suck, G.; Odendahl, M.; Nowakowska, P.; Seidl, C.; Wels, W.S.; Klingemann, H. G.; Tonn, T. Cancer Immunol. Immunother. 2016, 65, 485.

(27) Munn, D. H.; Bronte, V. Curr. Opin. Immunol. 2016, 39, 1.

(28) Zou, W.; Wolchok, J. D.; Chen, L. Sci. Transl. Med. 2016, 8, 328rv4.

(29) Schönfeld, K.; Sahm, C.; Zhang, C.; Naundorf, S.; Brendel, C.; Odendahl, M.; Nowakowska, P.; Bönig, H.; Köhl, U.; Kloess, S.; Köhler, S.; Holtgreve-Grez, H.; Jauch, A.; Schmidt, M.; Schubert, R.; Kühlcke, K.; Seifried, E.; Klingemann, H. G.; Rieger, M. A.; Tonn, T.; Grez, M.; Wels, W. S. Mol. Ther. 2015, 23, 330.

## Reference

1. Wang, W. et al. Chemoenzymatic synthesis of GDP-l-fucose and the Lewis X glycan derivatives. Proc. Natl. Acad. Sci. USA 106, 16096–16101 (2009).

2. Besanceney-Webler, C. et al. Increasing the Efficacy of Bioorthogonal Click Reactions for Bioconjugation: A Comparative Study. Angew. Chem. Int. Ed. 50, 8051–8056 (2011).

3. Chen, L. et al. Improved variants of SrtA for site-specific conjugation on antibodies and proteins with high efficiency. Scientific Reports 6, 31899 (2016).

4. Jeong, H.-J., Abhiraman, G.C., Story, C.M., Ingram, J.R. & Dougan, S.K. Generation of Ca2+-independent sortase A mutants with enhanced activity for protein and cell surface labeling. PLOS ONE 12, e0189068 (2017).

5. Rahim, M.K., Kota, R. & Haun, J.B. Enhancing Reactivity for Bioorthogonal Pretargeting by Unmasking Antibody-Conjugated trans-Cyclooctenes. Bioconjug. Chem. 26, 352–360 (2015).

6. Rondon, A. et al. Antibody PEGylation in bioorthogonal pretargeting with trans-cyclooctene/tetrazine cycloaddition: in vitro and in vivo evaluation in colorectal cancer models. Scientific Reports 7, 14918 (2017).

